# QBayMic: Quantum-coupled variational Bayes for clustering and feature selection in low-signal microbiome data

**DOI:** 10.64898/2026.09.14.751634

**Authors:** Tung Dang, Artem Lysenko, Tatsuhiko Tsunoda

## Abstract

Clustering microbiome samples into community types is central to cohort stratification and biomarker discovery, yet the resulting inference becomes unstable when the between-group signal is small compared with sampling noise: variational Bayes yields different partitions across initialisations, and common fixes do not solve the problem. Simple restarts are ineffective because the variational free energy is anti-correlated with clustering accuracy; deterministic annealing collapses to the same solution as greedy ascent, with the operator staying diagonal at every temperature; and parallel-tempering replicas remain too similar to permit configuration exchanges. We propose QBayMic, which replaces the assignment step of a Dirichlet-multinomial mixture with sparse variable selection via a quantum Gibbs state under an annealed Hamiltonian, coupling competing assignments through a transverse-field term that cannot be reproduced by temperature scaling alone. We present two gate-based circuit designs for this step, evaluating on noiseless qubit-register simulations, and we derive a signal fraction *σ*, computable prior to clustering, that predicts the expected strength of quantum coupling. With matched compute in the predicted regime, the three classical methods recovered the reference partition (ARI > 0.4) in 0/100 seeds, while QBayMic recovered it in 47–64/100; when the number of clusters exceeded three, only QBayMic recovered the correct cluster count. For a soil pH dataset, the diagnostic indicates a narrow separation margin; for a human-derived dataset tuned into the predicted band via controlled dilution, classical methods recovered the cluster count in 0/100 seeds, compared with 61–76% for QBayMic. The implementation is publicly available at https://github.com/tungtokyo1108/QBayMic.

## 1. Introduction

High-dimensional microbiome sequencing profiles thousands of taxa across multiple samples. Inferring the latent community structure supports disease profiling, cohort stratification, and therapeutic discovery [1, 2, 3]. Two targets are central: clustering samples into community types and identifying sparse taxa that distinguish them as candidate biomarkers. Probabilistic mixture models are well-suited for these settings because they directly model count data. In particular, a sample’s total count primarily reflects the sequencing depth, and replicate samples within a given community type often exhibit variability exceeding that permitted by a multinomial model. Introducing a Dirichlet prior on the multinomial parameter yields a Dirichlet–multinomial distribution that accommodates overdispersion; mixtures of such components represent latent subgroups, and a stick-breaking prior enables the number of components (*K*) to be inferred [4, 5]. The framework can be augmented with sparse variable selection to address application-specific objectives [6]. Empirically, only a minority of measured taxa tends to differ systematically between community types; retaining a shared background of non-differential taxa allows this uninformative majority to dominate the likelihood and obscure the discriminative signals. Accordingly, a per-taxon inclusion indicator estimated jointly with clustering can restrict the inference to an informative subset. Here, we adopt the Dirichlet–multinomial mixture with stochastic variational variable selection (DMM-SVVS) [6] and modify only the procedure used to compute the cluster assignments. The key challenge is inference: the count matrices are sparse and strongly zero-inflated, and the discriminative signal is weak relative to the compositional noise [7, 8], resulting in a highly multimodal objective landscape.

Mean-field variational inference is a standard scalable approach that updates cluster assignment probabilities via a softmax of candidate energies [9, 10, 11]. In low-signal metagenomic settings, coordinate-ascent optimization frequently converges to spurious local optima determined by stochastic initialization [12, 13]. Three established mitigation strategies are commonly employed. First, one can perform multiple random restarts and retain the solution that achieves the minimal variational free energy [14, 15, 16]. Second, the optimization landscape can be modified using deterministic annealing variational Bayes (DAVB), which introduces an elevated temperature to maintain diffuse assignments and subsequently cools according to a schedule, yielding sharper partitions only as the temperature decreases [17, 18, 19]. Third, the posterior can be explored using parallel tempering (PT), which simulates multiple replicas across an inverse-temperature ladder and proposes swaps between adjacent temperatures, enabling high-temperature replicas to traverse modes and transmit configurations to low-temperature replicas that target the posterior [20, 21]. These annealing and replica-exchange methods are strong baselines in related domains, including speech recognition [22], signal processing [23], structural bioinformatics [24, 25], and protein folding [26]. However, none of these approaches transfer effectively to the present regime, and our empirical evaluations clarify the failure mode in each case. With restarts, the issue is model selection: variational free energy is anti-correlated with clustering accuracy; thus, choosing by free energy prefers worse partitions. With DAVB, all temperatures converge to the same greedy fixed point because the assignment update remains diagonal; thus, annealing changes the path but not the solution. With PT, we cannot construct an effective temperature ladder: the swap acceptance stays around 55– 60% independent of the maximum temperature, above the 20–40% range, suggesting distinct replica states. Therefore, replicas remain in one basin and exchange insufficiently diverse configurations.

Quantum annealing provides an alternative method for exploring rugged energy landscapes [27, 28, 29, 30]. In quantum annealing variational Bayes (QAVB), the assignment update is reformulated in an operator-theoretic form [31, 32, 33]: assignment energies become diagonal operator entries, a mixing (driver) term introduces off-diagonal couplings between competing assignments, and responsibilities are computed from the Gibbs state of an annealed Hamiltonian. When the mixing term vanishes, QAVB reduces to the classical variational update, making it a one-parameter deformation of standard variational inference. The mixer enables the probability mass to move among competing assignment behaviors that are inaccessible to purely diagonal updates at any temperature. Despite this motivation, implementations have remained classical: updates avoid explicit density matrix calculations and differ only modestly from deterministic annealed variational Bayes [31]. Although the qubit-level resource requirements have been stated ⌈log*K*⌉ qubits and *O*(*K*) operations per update step [33], a gate-level circuit has not been provided for the single-qudit cyclic-shift operator that provides the mixing interaction. Empirical studies mostly compare QAVB to DAVB rather than replica-exchange baselines, leaving the gains untested against strong, classical alternatives. To our knowledge, QAVB has not been applied to high-dimensional, sparse, zero-inflated biological count data, and current approaches do not offer a principled way to predict which datasets will benefit from quantum approaches.

We address these gaps with QBayMic (**Q**uantum **Bay**esian **Mic**robiome) and make four contributions. First, we propose quantum-annealed inference to replace the cluster assignment step of DMM-SVVS with the preparation of a Gibbs state under an annealed Hamiltonian and leave the generative model, sparse selection layer, and all remaining update equations unchanged. This substitution recovers the community structure that the classical inference of the same model does not find. Moreover, we construct a quantum circuit that QAVB argued to be feasible but did not build. Sample assignments are encoded in registers of *n*_*q*_=⌈log*K*⌉ qubits with transverse-field mixing; therefore, the register width depends only on the number of clusters and is independent of the sample *N* and microbial taxa *S*.

Second, QAVB does not provide a circuit implementation of its cyclic-shift mixing operator; as a result, it must be executed classically, and the exact procedure, which we keep only for reference, constructs the complete 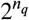× 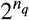 operator. We thus present two circuit-based implementations for preparing the annealed thermal state, each with a distinct resource trade-off. A variational quantum thermalizer (VQT) [34] generates the state on *n*_*q*_ qubits without ancillas, with a per-step cost that scales linearly with the number of circuit parameters. We choose this as the practical route for structural reasons: its classical latent ensemble makes the entropy contribution to the free energy exactly evaluable classically, eliminating the truly costly component of thermal-state preparation. Variational quantum imaginary-time evolution (VarQITE) [35, 36] produces the same state on a 2*n*_*q*_+1 qubit system–ancilla register via the dynamical projection approach, using McLachlan’s variational principle, with a per-step cost that is quadratic in the parameter count plus the cost of solving a linear system.

Third, we decompose the free-energy barrier, which coincides with the agglomerative information-bottleneck merge cost [37], into the population signal and the finite-sample noise. The resulting signal fraction *σ* is computable from the data before clustering and predicts where the quantum approach is beneficial. Across our sweep, the advantage is confined to *σ* between approximately 0.29 and 0.43. The decomposition also predicts that the height of the barrier does not govern difficulty, since a merge cost is large precisely when two groups are well separated, and the sweep confirms this. Controlling for *σ*, the partial correlation between success and barrier height is −0.32 and is not distinguishable from zero, whereas controlling for barrier height the partial correlation with *σ* remains +0.62.

Fourth, we assessed the synthetic dataset at varying difficulty levels and two real-world datasets using two endpoints: whether the inferred partition agrees with the reference labels (ARI exceeding 0.4) and whether the method recovers the correct number of clusters. Under comparable compute (*σ* ≈ 0.29), QBayMic recovered the correct partition in 47 to 64 out of 100 seeds on the benchmark cell dataset and recovered the group counts in 82 to 89% of seeds, whereas greedy variational Bayes, DAVB, and PT failed to recover either endpoint for any seed. For a continental soil pH dataset [38], the diagnostic yields *σ* = 0.60, implying an expected ARI advantage of about +0.01 to +0.03 over the best-performing classical method. For a cohort derived from human oral data [39] and diluted to match the predicted difficulty, the quantum methods recovered the cluster number in 61 to 76% of seeds, compared with 0 for all three classical baselines.

## 2. Materials and methods

### 2.1. Overview

QBayMic confines quantum computation to the one step where classical variational inference fails: the assignment of samples to community types, and the model of the priors and every other update remains classical (Fig. 1). Rather than attempting to encode a high-dimensional microbiome count matrix onto quantum hardware, each sample receives its own register of *n*_*q*_ = ⌈log *K*⌉ qubits encoding its *K* candidate cluster labels. Therefore, the quantum resource is fixed by the number of clusters and is independent of the number of samples *N* and the number of taxa *S*. The *N* registers are mutually independent and are evaluated in parallel.

**Figure 1:**
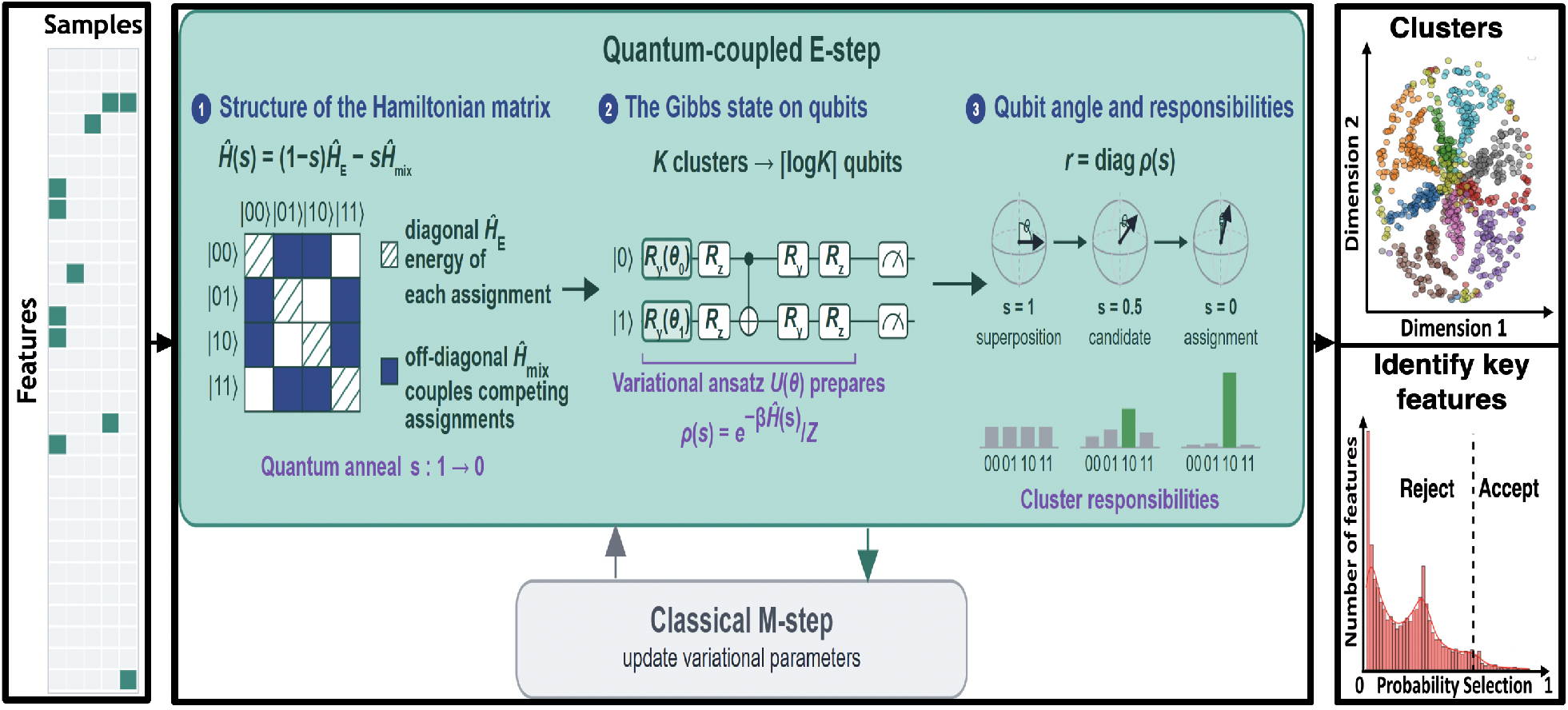
Schematic of the QBayMic model. (1) Structure of the operator. For a single sample, the annealed operator *Ĥ*(*s*) = (1 − *s*) *Ĥ*_E_ + *s*Γ *Ĥ*_mix_ is constructed from the current variational parameters. Its diagonal holds the assignment energy *Ĥ*_E_ of each candidate cluster label and reproduces the classical update independently. The off-diagonal entries come from the transverse-field mixer *Ĥ*_mix_and couple competing assignments, for which the classical update has no counterpart. The schedule runs *s* : 1 → 0, from mixer-dominated to energy-dominated state. **(2) Preparing the state on qubits**. Each sample’s assignment posterior is carried out on *n*_*q*_ = ⌈log *K*⌉ qubits. A parameterized ansatz *U* (*θ*) of single-qubit *R*_*y*_ and *R*_*z*_ rotations interleaved with CNOT entanglers rotates a classical mixture over basis states into the Gibbs state *ρ*(*s*) = *e*^−*β Ĥ*(*s*)^/*Z*. **(3) Readout**. Measuring in the computational basis gives the cluster responsibilities as the thermal populations *r* = diag *ρ*(*s*). The Bloch spheres and histograms show one sample across the schedule: near uniform at *s*=1, still holding candidates at *s*=0.5, resolved at *s*=0. **Classical M-step**. The responsibilities are passed to a conventional variational update of all remaining parameters, including the sparse feature selection indicators; the E and M steps alternate to convergence. The converged model returns the recovered community structure and the posterior feature selection probabilities.

Within the quantum-coupled E-step, the per-assignment energies that the classical update would pass to a soft-max are placed on the diagonal of an operator *Ĥ*_E_, and a transverse-field mixer *Ĥ*_mix_ supplies off-diagonal entries coupling competing assignments [40, 41]. These terms are combined under an annealing schedule that interpolates from a mixer-dominated regime to an energy-dominated regime. The departure from classical approaches is therefore structural rather than merely computational: greedy variational Bayes, DAVB, and PT effectively exponentiate a purely diagonal operator, and thus do not contain off-diagonal components at any computational budget. Consequently, they cannot directly transfer probability mass between competing assignments: temperature modulation only sharpens or flattens pre-existing weights without inducing transitions among assignments, and replica exchange moves entire configurations between replicas without generating within-replica coupling between alternatives. By contrast, the off-diagonal elements of *Ĥ*_mix_ directly couple different assignments, enabling transfer and producing the behavior in Figure 1: probability mass initially spread across many assignments early in the schedule gradually concentrates on a single assignment. Classically reproducing this requires the full operator, not just its diagonal, and constructing its dense exponential needs 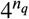 memory. For the register sizes used here this is still inexpensive, so we compute it exactly and use it to validate the circuit preparations.

To prepare the target thermal Gibbs state *ρ*(*s*) = *e*^−*β Ĥ*(*s*)^/*Z* without exponentially scaling classical matrix operations, QBayMic employs a parameterized variational quantum circuit ansatz *U*(*θ*) [34, 35, 36]. As illustrated in Figure 1, the circuit applies interleaved single-qubit rotations (*R*_*y*_, *R*_*z*_) and entangling CNOT gates to rotate a classical mixture over the computational basis states into the target thermal state, enabling an exact classical entropy evaluation at zero quantum measurement cost. Measuring the prepared density matrix in the computational basis extracts the diagonal thermal populations as cluster responsibilities *r*_*ik*_. These responsibilities are passed directly to a conventional classical M-step, which executes sparse variable selection to identify discriminating biomarker taxa and updates the model hyperparameters.

### 2.2. Generative model

We are given a microbiome count matrix 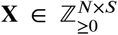 recording abundance of *S* taxa across *N* samples. Microbiome counts are sparse, compositional, and heavily zero-inflated, and the discriminating signal is usually carried by a small number of taxa. Therefore, we adopt the Dirichlet– multinomial mixture with stochastic variational variable selection (DMM–SVVS) [6], the established generative model for microbiome counts [4].

Sample *i* carries a latent group label *z*_*i*_ ∈ {1, …, *K*} drawn from Categorical(π), with a truncated stick-breaking prior on π so that the number of occupied groups is inferred instead of fixed; *K*_max_ denotes the truncation level. Conditional on *z*_*i*_ = *k*, the counts follow a Dirichlet–multinomial with group concentration *ϕ*_*k*_. A per-taxon Bernoulli indicator *γ*_*s*_ splits taxa into an informative set, which discriminates groups, and a shared background, which does not. This selection layer adapts the model to the high-dimensional sparse regime, where only a small fraction of taxa carry the group signal. Supplementary Material Appendix B provides the full hierarchical and selection priors.

Inference targets the joint posterior of ({*z*_*i*_}, π, {*ϕ*_*k*_}, {*γ*_*s*_}). Mean-field variational inference [10] approximates the posterior by a factorized distribution *q* and maximizes the evidence lower bound (ELBO). The assignment update is a softmax over a per-sample assignment energy,

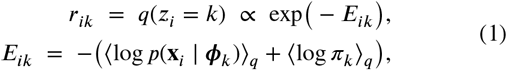

with ⟨⋅⟩_*q*_ being the expectation under the current *q*. The greedy coordinate ascent iterates Eq. (1) together with complementary parameter updates until it reaches a local optimum.

Two properties of this update determine what follows. First, it is strictly local: a sample moves only to the currently best-looking group, so the algorithm falls into the basin favored by initialization and cannot escape. The challenge is a rugged objective, not limited model capacity. Second, and more importantly in our experiments, the objective is a poor guide. In the low-signal reference setting, variational free energy anti-correlates with clustering accuracy (Pearson *r* ≈ −0.55 over 30 seeds). The global free-energy optimum is a noise-driven partition, while the biologically correct clustering is a higher-free-energy local optimum. Greedy inference can reach it, but selecting runs by free energy favors the spurious solution.

### 2.3. The quantum-coupled approach

#### 2.3.1. Construction

Following the quantum annealing variational Bayes (QAVB) construction [32, 33], QBayMic replaces the softmax assignment step in Eq. (1) with a quantity computed from quantum thermal states. For sample *i*, we represent the *K* candidate assignments using *n*_*q*_ = ⌈log *K*⌉ qubits, such that the basis state |*k⟩* represents the hypothesis *z*_*i*_ = *k*. From the same per-assignment energies *E*_*ik*_ that define the classical responsibilities, we build a diagonal energy operator 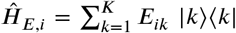. We separately introduce a mixing operator 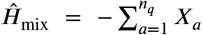, where *X*_*a*_ is the Pauli-*X* operator on the qubit *a*. Its role is to supply off-diagonal entries, that is, direct couplings between different candidate assignments, which the purely diagonal *Ĥ* _*E,i*_ does not have.

The two operators are combined along a schedule parameter *s* that runs from 1 to 0 during inference as follows:

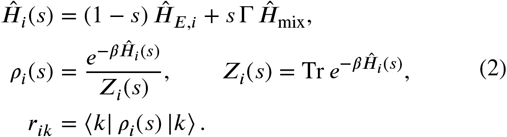

Here, *ρ*_*i*_(*s*) is a Gibbs (thermal) state at inverse temperature *β*, and the responsibilities are its diagonal entries, that is, the probability assigned to each candidate. Because *Ĥ* _*i*_(*s*) acts on a separate register for each sample, the joint state factorizes as *ρ* = ⨂ _*i*_ *ρ*_*i*_ (*s*): the register is only ⌈log *K*⌉ qubits wide, and the cost scales with the number of samples, not with the register size. For the mixing operator, we use the standard multi-qubit transverse field − ∑_*a*_ *X*_*a*_ instead of the single-qudit cyclic-shift mixer in the original QAVB formulation [32, 33]. Two reasons: the transverse field is the canonical quantum-annealing mixer [40, 41], and setting *Ĥ* _mix_=0 then gives an exact classical annealing limit. Explanations are provided in Supplementary Material Appendix E.

#### 2.3.2. Why this can help

The two ends of the schedule are easily described. At *s*=1, the operator is a pure mixer, and its thermal state spreads uniformly over all assignments. At *s*=0, it is purely diagonal, and *ρ*_*i*_(0) reproduces exactly the classical responsibilities of Eq. (1). The method is a one-parameter family interpolating between a fully undecided assignment and the standard greedy update, with the classical E-step recovered as the *s*=0 member. In between, the mixing term gives nonzero couplings ⟨*k*| *Ĥ*_*i*_ (*s*) | *k*^′^ ⟩ between competing assignments; thus, the probability mass can be redistributed among low-energy candidates as the schedule progresses. The classical update, being strictly diagonal, has no analogous move; it can only descend within whichever basin it already occupies. Informally, the method keeps competing assignments in play until the schedule resolves them, instead of committing each sample to whichever group currently looks best.

The mixing term acts as a fluctuation that smooths the assignment landscape, playing a role analogous to the thermal fluctuation of deterministic annealing, but is distinct from it. This distinction is exactly what our control isolates: a purely classical anneal following the identical schedule with *Ĥ*_mix_=0stays diagonal throughout, so any advantage of QBayMic over that control is attributable to the off-diagonal coupling and not to annealing alone.

### 2.4. Quantum circuit preparation of the Gibbs state

#### 2.4.1. Variational quantum thermalizer

The assignment step of Eq. (2) requires the Gibbs state 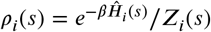 on the *n*_*q*_ -qubit register. Obtaining it directly means forming *Ĥ*_*i*_ (*s*) as an explicit 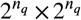 matrix and exponentiating it; thus, the memory grows as 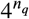, and the approach is limited to a handful of qubits. We prepare the state with a parameterized circuit instead, which does not require the construction of the matrix.

One obstacle must be addressed first, which shapes the construction. A circuit applied to a fixed input produces a pure state, that is, a single vector. The thermal state at a finite temperature is a probabilistic mixture of several states. Therefore, a circuit alone cannot produce it. The variational quantum thermalizer (VQT) [34, 42] resolves this by supplying the mixedness classically and rotation quantum mechanically.

Write |*x*⟩ for the computational basis state labelled by the bitstring 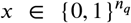 the register has 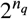 of them, and the first *K* represent the candidate assignments, for which we write |*k*⟩. The VQT samples *x* from a classical distribution *p*_*η*_ (*x*) |*x*⟩, prepares, and applies a parameterized circuit *U*(*θ*):

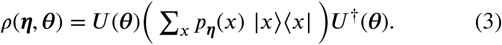

The bracketed term is a classical mixture over basis states, diagonal and manifestly mixed, and *U*(*θ*) rotates it into position. We take *p*_*η*_ to factorize across qubits, so *η* holds one Bernoulli parameter per qubit, and *θ* holds the circuit angles. Both are classical parameters and are trained together.

Training minimises the variational free energy

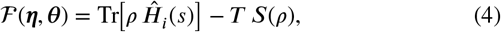

whose minimizer over all states is the target *ρ*_*i*_(*s*), so driving ℱ down drives Eq. (3) toward the desired state. The construction fits this problem for a reason worth making explicit. Free-energy minimisation requires the von Neumann entropy *S*(*ρ*), and estimating the entropy of a state held in a quantum register is normally the expensive part of thermal-state preparation. Here it costs nothing. *U*(*θ*) is unitary, so it leaves the eigenvalues of whatever it acts on, unchanged. The eigenvalues of *ρ*(*η, θ*) are therefore exactly the classical weights *p*_*η*_(*x*), and

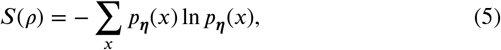

The Shannon entropy of a distribution is already well-known. It is computed exactly on the classical side at no quantum cost. The gradient of ℱ then costs *O*(|θ|) per step, with no metric to construct or invert. Once ℱ is minimised, the responsibilities are read off as the diagonal of the prepared state in the assignment basis, *r*_*ik*_= ⟨ |*ρ* (*η* ^*⋆*,^ *θ*^*⋆*^)|*k* ⟩. The register stays at *n*_*q*_=[log_2_ *K*] qubits throughout, no ancilla is required, and no step of the procedure forms a 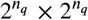 matrix. Mathematical explanations are provided in Supplementary Material Appendix C3.

#### 2.4.2. Variational quantum imaginary-time evolution

Variational quantum imaginary-time evolution (VarQITE) [35, 36] reaches the target using a different route. Under imaginary-time evolution, an initial state relaxes toward the thermal state as *β* grows; however, the exact trajectory generally leaves the set of states that a fixed circuit can represent, so each step is projected back onto it. McLachlan’s variational principle selects the parameter velocity 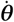 that best matches the true evolution, obtained by solving

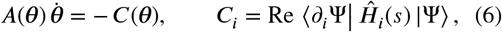

where |Ψ(*θ*) ⟩ is a parameterized purification of a 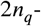 qubit system–ancilla register whose reduced state carries the target populations, and *A* is the quantum Fisher information metric, which measures how far the state moves per unit change in each parameter. Assembling *A* costs *O* (|*θ*|^2^) derivative evaluations per step plus a linear solve, making the VarQITE the most expensive of the three. Its value here is that it reaches the target using a mechanism that shares nothing with Eq. (4), so the agreement between the two is evidence of the target rather than a single optimizer. Mathematical explanations are provided in Supplementary Material Appendix C4.

#### 2.4.3. Exact diagonalisation

Exact diagonalization (ED) computes the target directly without variational approximation and provides the ground truth for the other two. Classically, *Ĥ*_*i*_(*s*) is formed as an explicit matrix and diagonalised, giving

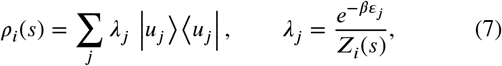

Where *ε*_*j*_ and |*u*_*j*_ ⟩ are the eigenvalues and eigenvectors of *Ĥ*_*i*_(*s*), and *λ*_*j*_ is the Boltzmann weight of eigenstate *j*. These *λ*_*j*_ are the populations that VQT approximates with its classical weights *P*_*η*_(*x*). The eigenpairs are then assembled into the purification 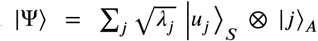. the same object VarQITE approaches variationally, which is constructed exactly. The square roots are what make the ancilla trace return the target: discarding *A* leaves exactly Eq. (7). We load | Ψ⟩ into the PennyLane state-vector simulator and trace over the ancilla, giving *r*_*ik*_ = ⟨*k* |*ρ*_*S*_ |*k*⟩ with *ρ*_*S*_ = Tr_*A*_ |Ψ⟩⟨Ψ|.

The last step does not change the value; it simply enforces that all three approaches employ the same readout code, so any observed discrepancies arise from the quality of state preparation rather than from measurement. ED is a classical, non-scalable method because its memory requirement scales as 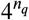. We use it as a benchmark to check the other two approaches. This should not be interpreted as a strict performance ceiling, and that nuance matters for the discussion ahead. By definition, any circuit result that differs from ED is less faithful to the intended Gibbs state; nonetheless, at the cluster-count endpoint, such a difference is not purely detrimental. Depending on the regime, circuit-based preparations identify the correct number of clusters more often than the ED method (Section 4.1.3). Preparation noise changes the trajectory of the inference rather than worsening it in a strictly monotonic fashion, meaning ED constrains fidelity to the target state rather than guaranteeing the fit outcome.

#### 2.4.4. Algorithm

Algorithm 1 provides an outer inference loop. This is standard variational inference for DMM–SVVS, except at line 6, where the softmax assignment update is replaced by a call to Algorithm 2. The M-step remains unchanged; therefore, the comparison with greedy variational inference in Section 3.3 differs in exactly one component.

Algorithm 2 presents the quantum-coupled E-step. Two features of it matter for the cost. First, the annealed operator is held in the Pauli basis rather than as a matrix: the diagonal energy block is expanded in Walsh terms, and the mixer is a sum of single-qubit *X* operators, so no step of the procedure allocates a 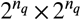 array. Second, the circuit and latent parameters are carried over between outer iterations (line 11). Successive schedule points differ only slightly, so the warm-started inner loop converges in a few steps, and *M* can be small.

The schedule defines *s*_*t*_ = *s*_0_ max(1 − *t*/*τ*_1_, 0), which makes *s* drop to zero precisely at iteration *τ*_1_; after that, the E-step reduces to the usual variational update. Moreover, the convergence check is prevented from triggering until iteration *τ*_2_ > *τ*_1_, ensuring that each run carries out at least *τ*_2_ − *τ*_1_ iterations of the purely classical update before it can terminate. Consequently, the final output is always a fixed point of the standard variational update: the quantum-coupled stage influences which fixed point the algorithm converges to, but it does not change the definition of a valid solution. Further details on the parameter-setting annealing schedule and verification appear in Supplementary Material Appendix G.

Code listings split tasks: quantum operations prepare and measure circuit states, while a classical computer computes entropy (via Eq. (5)), free energy, and parameter updates. Each outer iteration uses *N* independent registers of *n*_*q*_ = ⌈log *K*⌉ qubits (one per sample). Supplementary Table S4 lists all settings for the generative model (truncated stick-breaking prior, concentration, selection prior), annealing schedule, circuit ansatz, and optimizer, including benchmark values and robustness-study ranges (Section 4.1). It also specifies *s*(*t*), *β*(*t*), mixer strength Γ, and the occupancy rule defining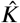.

##### Algorithm 1

QBayMic-VQT inference

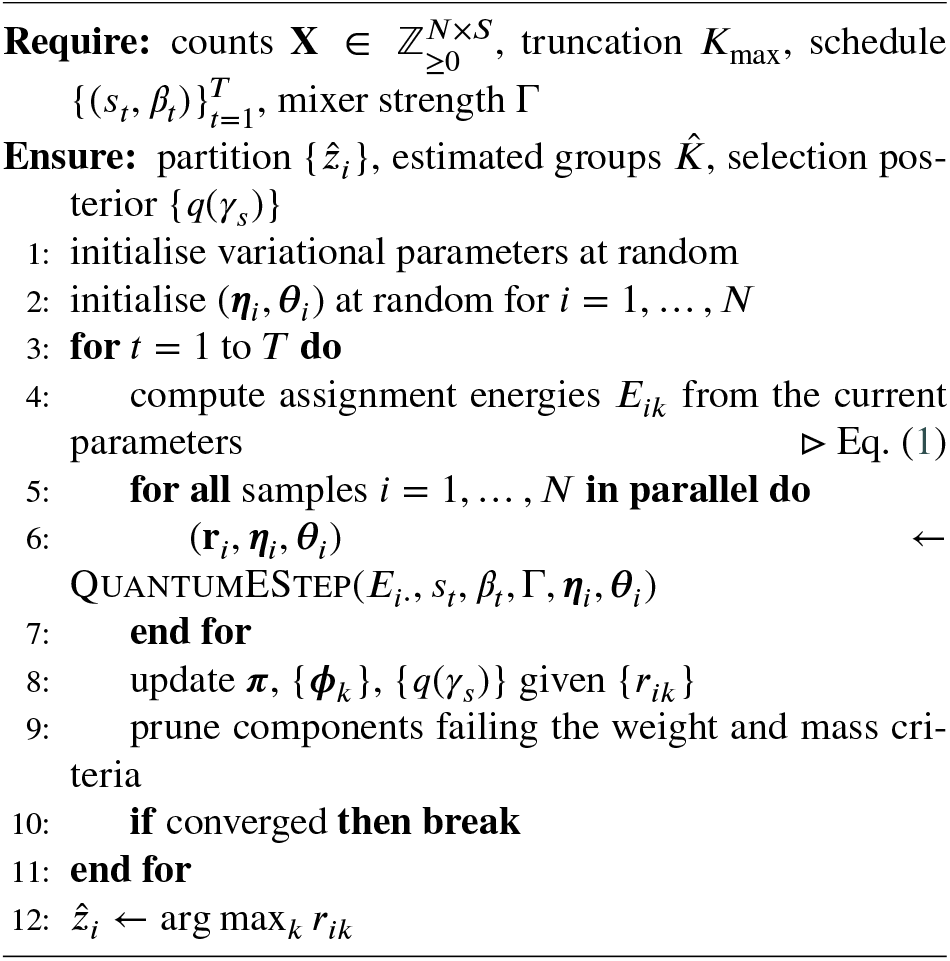

##### Algorithm 2

QuantumEStep: circuit preparation of the assignment posterior for one sample. Steps marked *(q)* run on the quantum register; the rest are classical.

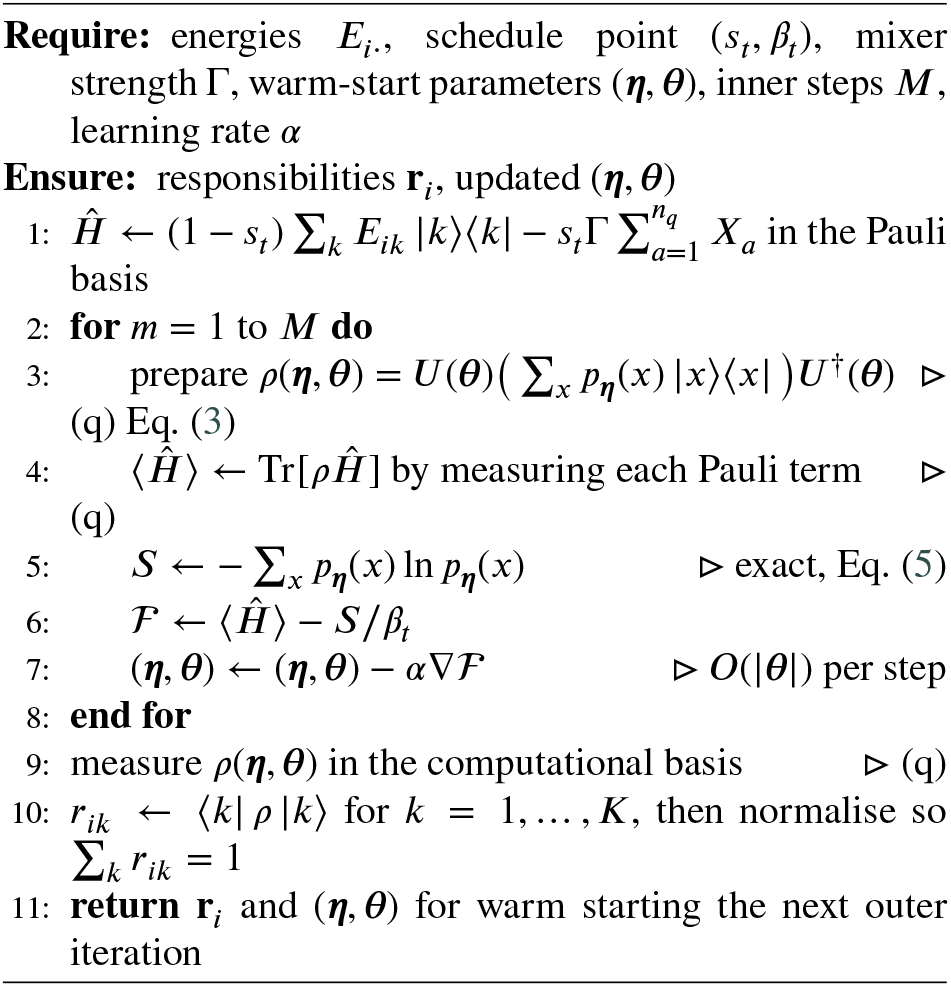

#### 2.4.5. Computational cost

Both variational procedures use a hardware-efficient brick-wall circuit: single-qubit ***R***_*y*_, ***R***_*z*_ rotations on each register qubit followed by a ladder/ring of CNOTs. They differ only in target register and depth: VQT uses the *n*_*q*_ -qubit system register at depth *d*, while VarQITE uses a 2*n*_*q*_+1-qubit system–ancilla register. Table 1 summarizes roles and costs.

**Table 1.**
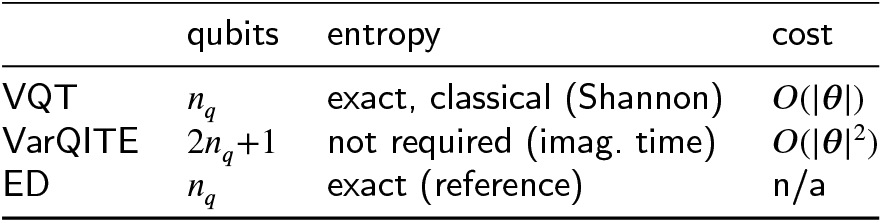
The three preparation procedures, with *n*_*q*_ = log *K*. Supplementary Material Appendix C5 and D give the per-group resource breakdown.

All three methods were simulated as exact, noiseless state vectors (Pennylane [43]), so reported effects reflect the inference algorithm rather than hardware noise. The Pauli decomposition reproduces the diagonal energy operator *Ĥ*_*E,i*_ to machine precision (max absolute error < 10−12), matching Eq. (2). At the schedule endpoint *S*=0, each method returns the classical responsibilities of Eq. (1) within 10−12, confirming the classical limit. On the clustering register, VQT and VarQITE match ED populations within integration tolerance; thus all three agree on the same state and hard assignments.

## 3. Experimental setup

### 3.1. Signal fraction *σ*

The size of the difference between the groups does not make clustering instances difficult. This is the extent to which the difference survives the finite sampling. We make this precise with a quantity and compute it for each instance in this study. Consider two candidate groups with fitted taxon profiles 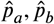 and post-zero-inflation total counts 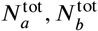. The energy cost of forcing them through a single centre, instead of keeping two, is

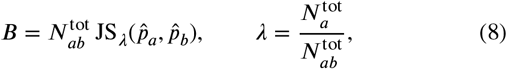

where JS_λ_ is the weighted Jensen-Shannon divergence [44]. It is the merge cost of the agglomerative Information Bottleneck [37] and is equivalent to the likelihood-ratio statistic for the homogeneity of two multinomial samples. Supplementary Material Appendix F provides the derivation in our notation because the decomposition below depends on this notation.

Recognizing *B* as a merge cost determines what it measures. The merge cost is high when two groups are well separated; therefore, a tall barrier indicates an easy instance. Barrier height therefore does not order inference difficulty; the composition of *B* does. Write 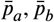 for the population profiles that the fitted 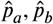 estimate. Splitting Eq. (8) at the population value gives

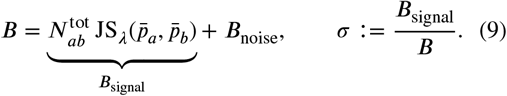

This split is definitional. What it separates is not: *B*_signal_ is the part of the barrier that reflects a real difference between populations, and *B*_noise_ is the part contributed by estimating high-dimensional sparse profiles from finitely many counts (Supplementary Material Appendix F2 and F3). Plug-in divergence estimators are biased upward in this regime; thus, *B*_noise_ is non-negative in expectation, and we verified that it is positive in every cell of the sweep. A low *σ* describes an instance whose barrier is mostly an artifact of estimation noise, and greedy inference freezes onto a spurious partition on these instances before the true split emerges.

A mechanism for this is available in the deterministic annealing literature, although we borrow it rather than derive it. In deterministic annealing, the correct split crystallises at a temperature set by the per-count signal *B*_signal_/*N*^tot^ [45, 17], so when that quantity is small, the split temperature falls below the point at which a descent has already been committed. We use this to motivate *σ*, not to prove anything about it. Our claim that *σ* predicts single-run reliability is empirical and established over the regime sweep in Section 4.

#### Computing *σ* in practice

For the synthetic data, *σ* is known exactly. The generator fixes the population profiles 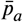, so 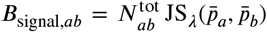 is the noise-free separation, and *B*_noise,*ab*_ follows by subtraction. Group membership is the generative label, so no partition is inferred and no estimator is involved.

Both grouping and population profiles must be supplied for real data. When an external grouping is available, as in the case of the cohort in Section 4.2, we use it directly. Otherwise, we obtain a preliminary partition from a single fast clustering of the relative abundance profiles (one greedy variational-Bayes pass), and *σ*_*ab*_ describes the separability of those fitted groups. We then estimate the sampling contribution to *B*_*ab*_ by resampling within groups, which yields comparisons between profiles drawn from the same distribution. The estimate must match the count budget it corrects: plug-in divergence bias scales as 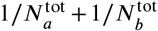, so a half-sized null overstates the inflation by roughly factor of two and drives *σ* toward zero. We therefore evaluate the null at several subsample sizes and extrapolate in inverse count, giving *B*_signal,*ab*_ = max(*B*_*ab*_ − *B*_noise,*ab*_, 0).

For *K* > 2 the barrier is not a single number but 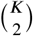 pairwise barriers, and difficulty is gated by the hardest 562 pair, so the reported quantity is *σ*_min_ = min_*ab*_ *σ*_*ab*_. In the 563 symmetric synthetic design, all pairs are equivalent and *σ*_min_= *σ* Each pair costs one Jensen–Shannon evaluation plus the resampled evaluations of the null on real data, so the diagnostic remains cheap relative to a single clustering run (Supplementary Material Appendix F5).

### 3.2. Data

Synthetic count matrices were generated with controlled separation, zero inflation, and informative taxon fractions [8]. Group profiles are drawn at three concentration levels (informative, cross-group, and background), library sizes are drawn from a negative binomial with mean 8000, and entries are independently zeroed with zero inflation probability. Two knobs set the difficulty of an instance: a separation parameter controlling the between-group signal and zero-inflation level. The barrier diagnostic in Section 3.1 maps every generated instance onto one difficulty axis, the signal fraction *σ*, and all synthetic results are reported accordingly. The main sweep is a 4 × 4 grid of separation ∈ {0.2, 0.3, 0.4, 0.5} and zero-inflation ∈ {0.6, 0.7, 0.8, 0.9}, which populates *σ* from the unrecoverable regime through the advantage band to the signal-dominated regime. The scaling study in Section 4.1.3 varies separation over the same four values at fixed zero-inflation 0.8, for each of *K* ∈ {3, 4, 5, 6}. One cell of the grid, with a separation 0.2 zero-inflation 0.80 and *σ* ≈ 0.29, serves as a reference configuration for the benchmark of Section 4.1.1 and for the barrier decomposition.

Continental soil pH (88-soils; *N* = 89 samples, *S* = 199 OTUs, *K* = 4 pH quantile bins) provides a natural gradient of overlapping communities [38]. Its measured signal fraction, *σ*_min_ = 0.60, places it in the signal-dominated regime; therefore, the prediction is a small margin rather than a qualitative separation. We also screened human labels from the Human Microbiome Project for a cohort inside the advantage band and found none: body sites are too well separated (*σ*_min_ > 0.85, so classical inference already succeeds) [39, 46, 47], and the disease labels we examined carry no recoverable community signal (*σ* ≈ 0, so no method succeeds) [48, 49, 50]. The intermediate regime appears to be uncommon among the natural labels available to us, which is worth reporting in Section 4.2.2.

### 3.3. Comparison of inference methods

To correctly attribute any advantage, we compared it with classical methods.

- **Greedy variational inference (VB)**. Eq. (2) at *S*=0; the standard approach and direct point of comparison.
- **Deterministic-annealing VB (DAVB)**. An identical temperature schedule with the mixing term removed [18, 19]. This is the decisive control that separates any benefit of annealing from any benefit of quantum coupling.
- **Parallel tempering (PT)**. We use replica exchange (PT) with an *M*-rung geometric inverse-temperature ladder whose cold replica (*β*=1) matches greedy VB, and Metropolis swaps between adjacent rungs [20, 21]. To make PT a fair baseline, we reseed hot replicas from fresh basins and gate swaps to match group count. PT is the strongest classical method for this multimodal inference. We tried to tune for the usual 20–40% swap-acceptance rate, but on this surface acceptance saturates at 55–60% regardless of the hot-end temperature because replicas fall into the same basin and overlap; we therefore report the closest ladder. This saturation suggests PT has no second basin to exchange into.

The clustering performance was assessed using the Adjusted Rand Index (ARI), which measures the agreement between the predicted cluster assignments *C* = {*C*_1_, …, *C*_*k*_} and true labels *T* = {*T*_1_, …, *T*_*m*_} while correcting for chance:

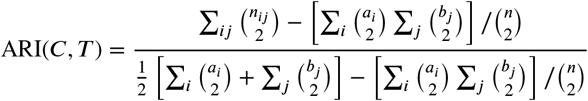

where *n*_*ij*_ is the number of samples in cluster *C*_*i*_ and true class *T*_*j*_, *a*_*i*_=Σ_*j*_ *n*_*ij*_, *b*_*j*_==Σ_*i*_ *n*_*ij*_, and is the total sample size. The ARI ranges from 0 (random agreement) to 1 (perfect agreement). The primary endpoint is single-run reliability: with hyperparameters fixed and only the random seed varying, we report the success probability *P* (ARI > *ϵ*) with Wilson 95% confidence intervals. The number of seeds was determined separately for each experiment. Over-specifying the truncation (*K*_max_ > *K*_true_) forces the model to estimate the number of groups 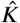, so we report a second and more difficult endpoint 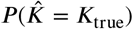. We also reported a parameter sensitivity analysis over a pre-registered grid, so that the results are shown across a reasonable parameter region and not at a single tuned point.

## 4. Results

### 4.1. Synthetic data experiments

#### 4.1.1. Single-run reliability at the reference cell

We began by benchmarking all methods on the low-signal reference cell (*σ* ≈ 0.29; *N* = 400 samples, *S* = 5000 OTUs, separation = 0.2, zero-inflation = 0.8, *K*_true_ = 3), while intentionally over-specifying the truncation at *K*_max_ = 4 (shared fixed parameters across methods: DP concentration = 3.2, selection prior = 0.7, prune threshold = 0.1, *τ*_1_ = 100, *τ*_2_ = 230) so that each method was required to infer the number of groups from the data. The random seed serves as the statistical unit: hyperparameters are fixed at a chosen operating point, ***R*** = 100 seeds are sampled and paired across methods, ensuring that all methods use the same initializations. In this setting, none of the three classical methods identified the correct structure in any run. Greedy VB, PT, and DAVB all achieved 0/100 successes at the threshold ARI > 0.4 (Fig. 2a and b), whereas the quantum-coupled E-step succeeded in 47 to 64 of 100 runs; correspondingly, the median ARI increased to 0.54 for the exact reference and to 0.39 for both quantum circuit preparations. Using PT as the strongest classical baseline for paired comparisons, the effect sizes were +0.47 ARI for ED and +0.35 for VarQITE and VQT, with bootstrap 95% intervals that excluded zero (Fig. 2d).

**Figure 2:**
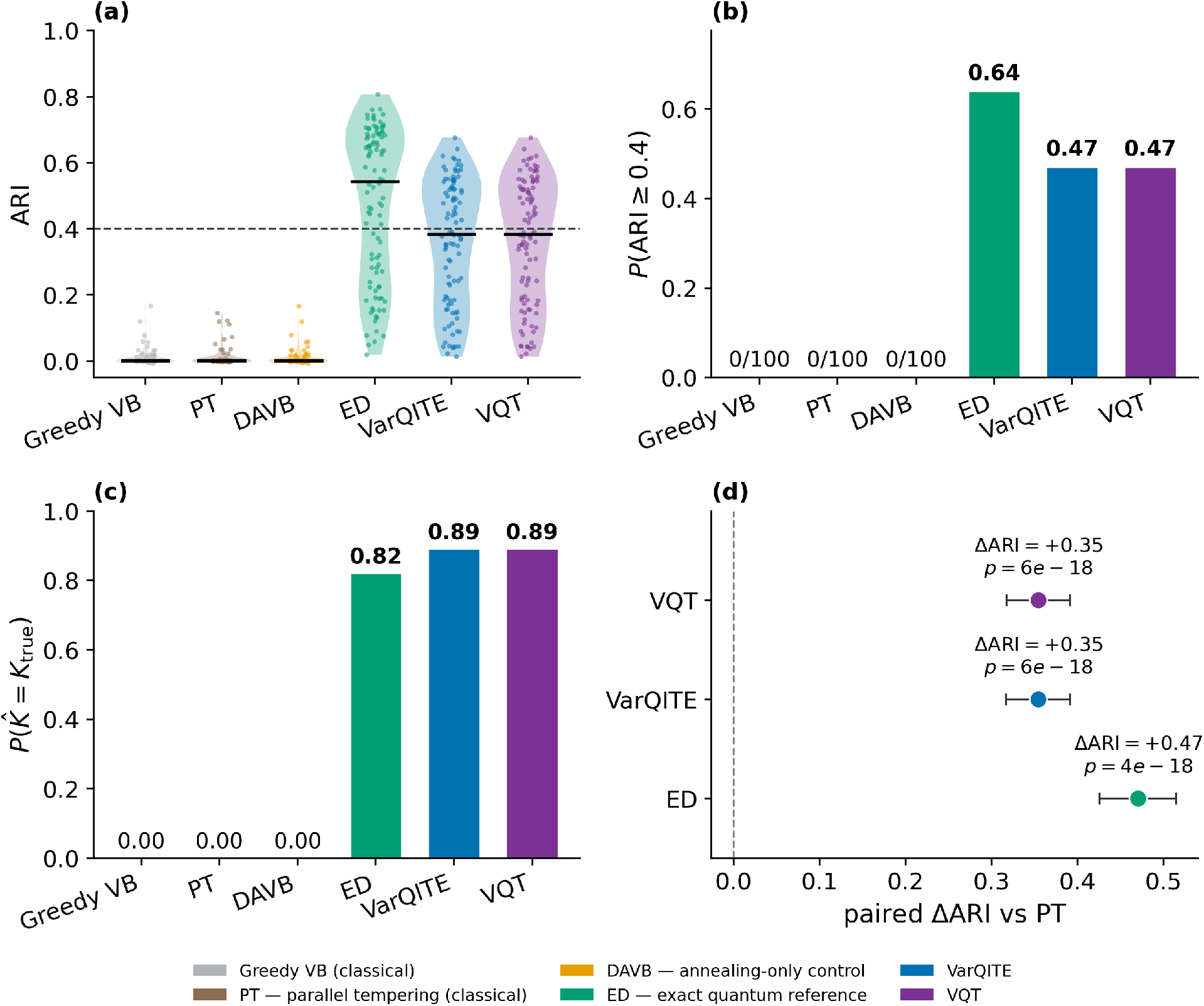
The quantum-coupled approach is reliable where classical inference fails. Benchmark at the low-signal reference cell (*σ* ≈ 0.29; 400 samples × 5000 OTUs, separation 0.2, zero-inflation 0.8; *K*_true_ = 3), *R* = 100 paired random seeds. (a) Per-seed adjusted Rand index (ARI); points are individual seeds, the horizontal bar is the median, and the dashed line marks the success threshold 0.4. (b) Single-run success probability *P* (ARI ≥ *ϵ* |*ϵ* = 0.4) in 100 seed. (c) Success probability of estimating the number of clusters 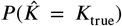 only the quantum E-step recovers the true cluster count. (d) Paired effect size relative to parallel tempering (ΔARI = ARIquantum − ARIPT); markers show the mean, error bars the 95% bootstrap confidence interval, and annotations the mean ΔARI with the paired Wilcoxon signed-rank *p*-value.

The key comparator for interpreting these results is DAVB, which uses the same temperature schedule but removes the mixing term and is otherwise an identical algorithm, including the same symmetry-breaking and internal heuristics. DAVB likewise scored 0/100; thus, annealing by itself does not solve this cell, and the performance gap relative to the quantum approaches is due to the off-diagonal coupling rather than the shared schedule. Parallel tempering was included as the strongest available classical approach for multimodal inference, and its failure was additionally revealed: swap acceptance could not be tuned into the typical 20 to 40% range and instead plateaued around 55% regardless of the hot-end temperature (Section 3.3).

The exact and quantum circuit preparations yielded different performances. ED achieved 0.64 (Wilson 95% [0.542,0.727]), compared with 0.47 (Wilson 95% [0.375, 0.567]) for both VQT and VarQITE. Thus, the deployable circuit approach reached approximately three-quarters of the reliability obtained by exact diagonalization, leaving an absolute gap of 0.17. Because these confidence intervals overlap, Figure 2b on its own makes the difference plausible but not definitive; however, a paired comparison across 100 hyperparameter draws (Supplementary Material Tables S1 and S2) clarifies the effect, showing a +0.07 median ARI advantage for ED over each scalable method, with intervals that exclude zero. In contrast, VarQITE and VQT matched each other within the measurement resolution at every reported endpoint. Because they optimized different objectives and did not share an optimizer, this agreement supports a conclusion about the target state rather than about either algorithm.

With over-specified truncation, the classical methods never recovered *K*_true_ = 3, while the quantum approaches did so in 82 to 89 out of 100 seeds (Fig. 2c). This criterion is more stringent than the ARI threshold, because a method may cluster accurately while still retaining an extra, spurious component. At the fixed operating point, the two quantum circuit preparations recovered *K* slightly more frequently than the exact method (0.89 versus 0.82). However, when averaging over random hyperparameters, the ranking flips and ED leads (0.35 versus 0.24 and 0.20; Supplementary Material Table S1). Both observations were reproducible. One possible explanation is that the variational circuit-preparation error provides a small smoothing effect that suppresses weak components at this specific configuration, an effect that does not persist when averaging across configurations.

Two additional checks addressed whether the findings were based on a single tuned setting or a single metric. Sampling 100 independent hyperparameter configurations per method (Supplementary Material TableS4) preserved the ordering: single-run success increased from 0.02 for greedy VB to between 0.23 and 0.42 for the quantum E-steps, and true-*K* recovery rose from 0.01 to between 0.20 and 0.35 (Supplementary Material Tables S1 and S2). The absolute rates are lower in that analysis because the search intentionally includes poor configurations; accordingly, the two analyses answer different questions (one about a selected operating point and the other about the hyperparameter prior) and should not be compared directly. Re-scoring the same runs using normalized mutual information (Supplementary Material Table S3) reproduced the same ordering and every pairwise conclusion. We nonetheless kept ARI as the primary metric because it is corrected for chance, which is important here: greedy VB is exactly 0 under ARI but exhibits a nonzero floor under NMI.

#### 4.1.2. The signal fraction predicts where the quantum advantage appears

We evaluated 16 cells across a grid of separation and zero inflation (400 samples × 5000 OTUs, *K*_true_ = 3, *K*_max_ = 4). For each cell, we computed *σ* prior to clustering and then executed each method using *R* = 15 paired random seeds under matched compute, with identical fixed settings throughout (DP concentration = 3.2, selection prior = 0.7, prune threshold = 0.1, *τ*_1_ = 100, *τ*_2_ = 230). For both quantum variants, single-run success increased monotonically with *σ* (Pearson *r* = +0.75 for ED and +0.75 for VQT; Fig. 3c), and the sweep separated into three regimes (Fig. 3a). When *σ* ≲ 0.24, no method recovered the latent structures. When *σ* ≳ 0.43, all methods succeeded and the particular E-step choice no longer affected outcomes. In the intermediate region, the quantum methods recovered the structure, whereas the classical methods did not, yielding advantage cells from roughly *σ* ≈ 0.29 to 0.43. We present this interval as an empirical observation rather than a sharp cutoff: the upper boundary lies between adjacent cells at *σ* = 0.419 and 0.427 which exhibit different classical performances, indicating dependence on the noise mechanism beyond what is summarized by *σ* alone (Section 3.1).

**Figure 3:**
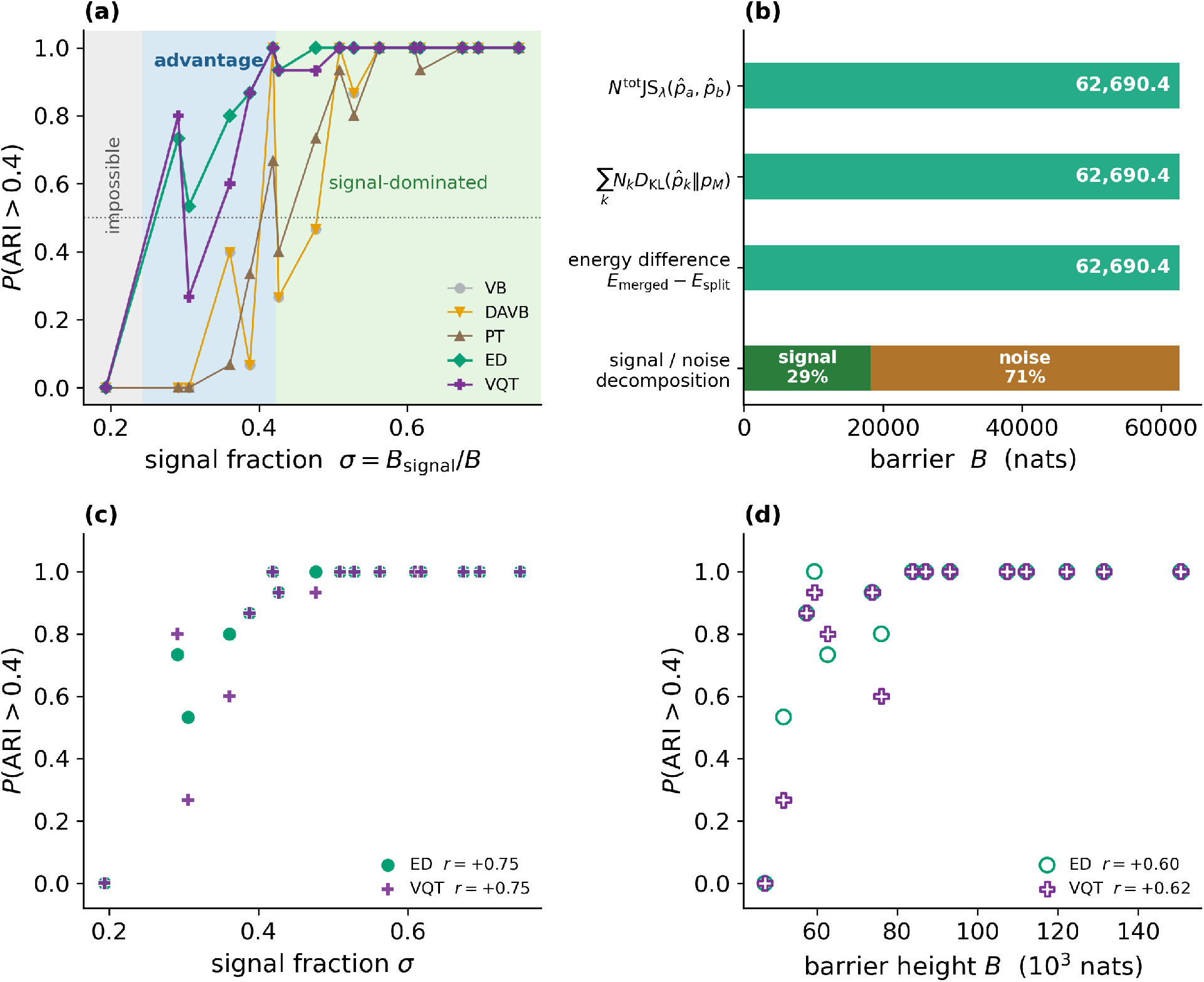
The signal fraction *σ* predicts when the quantum-coupled advantage appears. Regime sweep over 16 cells (separation × zero-inflation), 400 samples × 5000 OTUs, *K*_true_ = 3, *R* = 15 seeds; the signal fraction *σ* = *B*_signal_/*B* was computed for each cell. (a) Single-run success *P* (ARI ≥ *ϵ* | *ϵ* = 0.4) versus *σ*. Success collapses onto *σ*, defining three regimes separated by two onset thresholds: an **impossible** regime (*σ* ≲ 0.24); an **advantage** regime (0.24 ≲ *σ* ≲ 0.42) for QBayMic; and a **signal-dominated** regime (*σ* ≳ 0.42). (b) The barrier identity: *B* computed in three independent ways and decomposed into 29% signal / 71% noise at the reference cell (*σ* = 0.291). (c) QBayMic success collapses tightly onto *σ* (Pearson *r* = +0.75). (d) The same successes are not governed by barrier height *B*. The weaker positive correlation with *B* (*r* = +0.60) reflects co-variation of *B* with *σ* across the sweep.

The overall difficulty is governed by the barrier’s composition rather than its magnitude. For the reference cell, the barrier matched the reported precision across three independent calculations (Fig. 3b) and can be decomposed into 29% signal and 71% estimation noise. Although success appeared to correlate with barrier height *B* (*r* = +0.60 for ED and +0.62 for VQT; Fig. 3d), *B* and *σ* are themselves strongly coupled over the sweep (*r*_*σB*_ = +0.91). After controlling for *σ*, the partial correlation between success and *B* is −0.32 (ED) and −0.30 (VQT), neither distinguishable from zero; in contrast, controlling for *B* leaves the partial correlation with *σ* at +0.62 (Supplementary Material Table S5). Thus, we found no detectable independent contribution to the barrier height. This matches the interpretation of *B* as a merge cost: it is larger when clusters are more cleanly separated, so the highest barriers occur in the easiest instances. Given the collinearity and small sample size (*n* = 16 cells), we treated these partial correlation values as suggestive rather than definitive.

Deterministic annealing does not explain this result. DAVB matched greedy VB’s success in all 16 cells, yet followed different per-seed trajectories: intermediate agreement dropped to ARI ≈ 0.5 (up to 99% of samples differed mid-run), but both always converged to the same final partition (ARI(DAVB, VB) = 1.0). The schedule altered the path, not the endpoint. As shown in Supplementary Material Appendix F, a split crystallizes at a temperature set by its per-count barrier contribution, giving *T*_noise_/*T*_signal_ = *B*_noise_/*B*_signal_ = (1− *σ*)/*σ*. For *σ* < 0.43, the noise split appears at higher temperature, so cooling commits to the noise configuration before the true split is available. In the reference cell (*σ* ≈ 0.29), the ratio was 2.4, and it remained > 1 throughout the advantage regime (*σ* ≈ 0.29–0.43). Expanding the temperature range does not fix this; it only prolongs time in the noise-split region.

The quantum methods avoided this because the mixer does not act by lowering temperature. At *s* = 1 the assignment energies drop out, and the off-diagonal term *Ĥ*_mix_ directly couples competing assignments, so softening need not overcome an *O*(10^4^)-nat energy scale. Thus, the key difference from DAVB is the off-diagonal coupling, and DAVB’s failure reflects the landscape, not the chosen schedule.

#### 4.1.3. Scaling with the number of clusters

We repeated the sweep for *K*_true_ = 3 to 6 with an over-specified truncation *K*_max_ = *K*_true_ +1 (DP concentration 3.2, selection prior 0.7, prune threshold 0.1, *τ*_1_ = 100, *τ*_2_ = 230), so the cluster count must be inferred. We varied separation over {0.2, 0.3, 0.4, 0.5} at fixed *N* = 400, *S* = 5000, zero-inflation 0.8, with *R* = 15 paired seeds per cell (Fig. 4). For every *K*, results collapse onto *σ*: within each panel, all cells lie on a single success curve, and exact and circuit-based preparations agree. The changes with *K* are how classical methods sit relative to that curve: at *K* = 3 they recover structure across much of the range, but by *K* = 5 they are near zero, while quantum approaches still reach ≥ 0.80 at the upper end.

**Figure 4:**
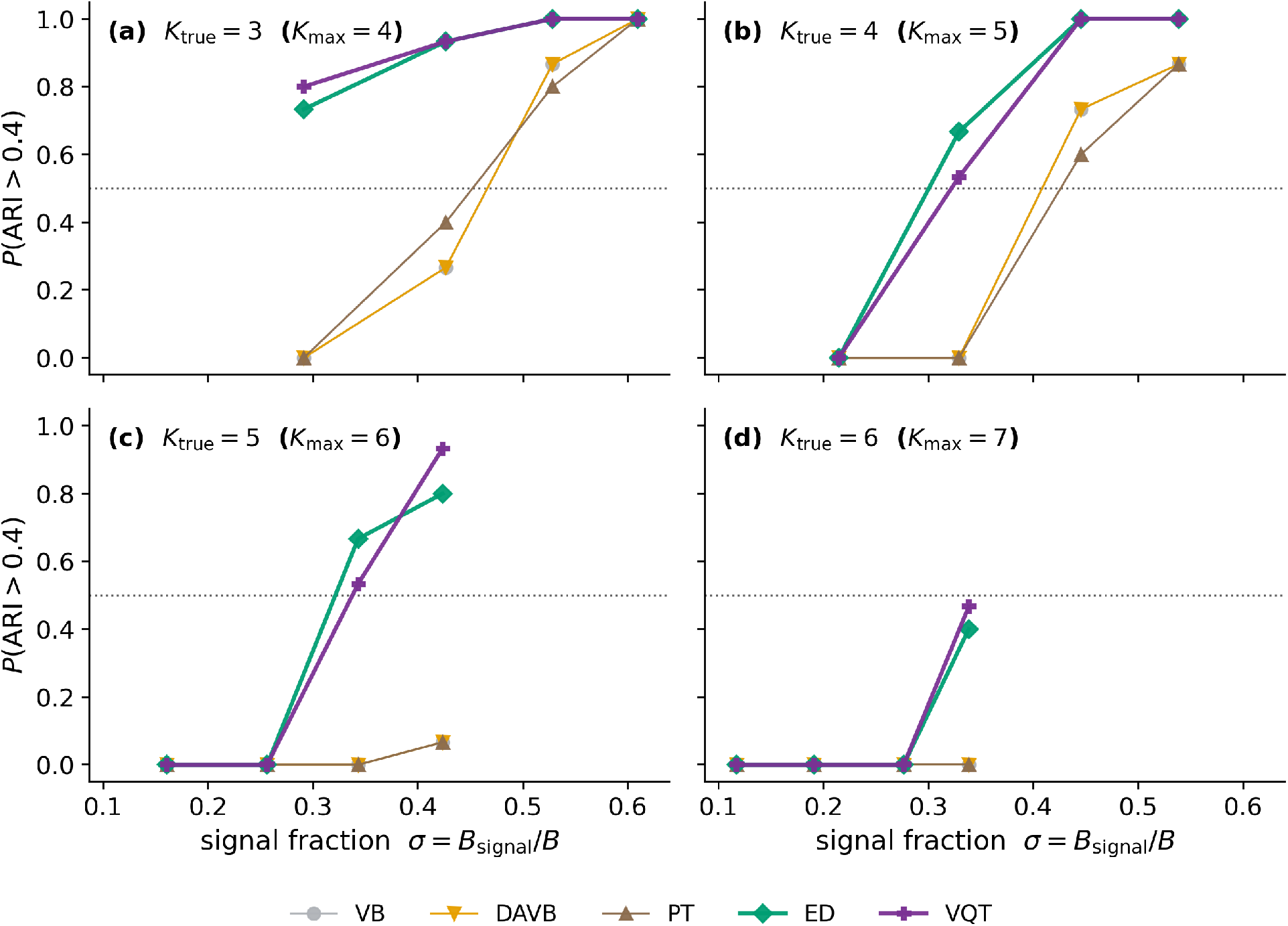
The quantum-coupled reliability advantage widens with the number of clusters. Single-run success *P* (ARI ≥ *ϵ*|*ϵ* = 0.4) versus data-computed signal fraction *σ* = *B*_signal_/*B* for *K*_true_ = 3 to 6 (panels a to d; *K*_max_ = *K*_true_ + 1), with *σ* swept via cluster separation ∈ {0.2, 0.3, 0.4, 0.5} at *N* = 400, *S* = 5000, zero-inflation = 0.8, *R* = 15 seeds.

In each panel, the four plotted points correspond to identical separation values. However, for any fixed separation, the resulting *σ* decreases as *K* increases, because the signal is distributed over a larger number of cluster pairs and consequently scales as 1/*K* (Supplementary Material Appendix F). Accordingly, in panel (a), the considered separations yield *σ* = 0.29–0.61, whereas in panel (d), they yield only 0.12–0.34. This implies that classical baselines operate in a more challenging regime at larger *K*. Nevertheless, the observed performance degradation is not solely attributable to increased difficulty; even at matched *σ*, performance decline persists. Specifically, at *σ* ≈ 0.42 (attained for both *K* = 3 and *K* = 5), DAVB and PT decrease from 0.27 and 0.40 to 0.00 and 0.07, respectively, whereas ED remains high (from 0.93 to 0.80) and VQT remains unchanged at 0.93 for both values of *K*.

The structure of the mixer provides a clear reason to anticipate this asymmetry. Increasing the number of clusters multiplies the ways a descent can end: pairwise merges, higher-order merges, and label confusions increase at least quadratically with *K* (Supplementary Material Appendix F). The classical update contains no off-diagonal entries anywhere in the schedule; as a result, it increases the number of detrimental spurious optima while lacking any mechanism to shift weight between assignments after committing. In contrast, the transverse field scales in the opposite direction. Operating on *n*_*q*_ = ⌈log *K*⌉ qubits, each term flips one qubit; consequently, every assignment is directly connected to *n*_*q*_ others and can reach any other in at most *n*_*q*_ flips: two flips for *K* = 3 and 4, and three for *K* = 5 and 6. Thus, as *K* grows, the assignment-space connectivity deteriorates only logarithmically, whereas the number of traps grows quadratically.

Recovering the correct number of clusters discriminated methods better than any accuracy threshold (Supplementary Material Fig. S1b) and was least sensitive to design choices, since *K*_true_ is either recovered or not. For *K*_*true*_ = 3, the best classical baseline recovered *K*_true_ in 0.20 of runs versus 0.58 for VQT. For *K*_*true*_ > 3, no classical method recovered the true value; at *K*_*true*_ = 4 recovery was 0.00 (Wilson 95% CI [0.00, 0.06]), while VQT reached 0.45 (Wilson 95% CI [0.33, 0.58]) and ED 0.25 (Wilson 95% CI [0.16, 0.37]), with CIs not overlapping the classical upper bound. The resulting separation is qualitative and directly affects whether practitioners infer the correct number of community types. This endpoint also favors the deployable circuit: VQT outperformed ED at both *K*_*true*_ = 3 and *K*_*true*_ = 4. The ordering matches the inversion in the reference cell (Fig. 2c) and reverses under random hyper-parameters (Supplementary Material Table S1), indicating a reproducible dependence on operating regime rather than transient variability.

The relative ranking of circuit-based state preparations versus ED in terms of *P* 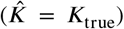 varies by condition, so we included further analyses. For synthetic cells with *K*_*true*_ = 3, ED tended to over-partition the data, identifying the correct number of clusters only 0.13–0.20 of the time (Supplementary Material Table S6). While this can be addressed in a setting by softening ED responsibilities using an explicit entropy-floor regulariser, the fix is both delicate and costly: the optimal regularisation strength depends on the dataset (the best floor differs across cells), and small mis-calibration can worsen recovery instead of improving it. By contrast, the VQT quantum circuit yielded a comparable softening effect without instance-specific tuning: even with a fixed variational circuit, performance matched or outperformed exact preparation (0.53 vs 0.13 at *σ* = 0.43). This pattern also appeared inherent rather than due to poor optimisation, since it persisted under greater ansatz depth and improved convergence. More broadly, circuit architecture provides an extra knob: swapping the transverse-field driver for a cyclic-shift mixer boosts recovery from 0.33 to 0.60 at *σ* = 0.53, surpassing even the best-tuned ED entropy-floor control. Overall, these findings indicate that mixer geometry and ansatz choices can provide a principled, tuning-free avenue for regularising model selection.

### 4.2. Real data experiments

#### 4.2.1. A natural cohort in the signal-dominated regime

We applied the method to the Lauber continental soil survey [38], clustering 89 samples across 199 OTUs into *K*_true_ = 4 balanced pH-quantile groups, with truncation *K*_max_ = 5 (fixed for all methods: DP concentration 0.77, selection prior 0.29, prune threshold 0.13, *τ*_1_ = 112, *τ*_2_ = 173) and *R* = 100 paired seeds. This dataset is a clean external test because the pH was measured (not defined by clustering), and pH-responsive taxa were known from the original study.

The diagnostic predicted only a modest benefit: the estimated signal fraction was *σ*_min_ ≈ 0.60 (Section 3.1), placing the data in the signal-dominated regime (Section 4.1.2), where all methods typically recover the pH structure, and quantum coupling should yield only a small gain. Single-run success *P* (ARI > 0.4) ranged from 0.89 (greedy VB) to 0.98 (ED) (Fig. 5b), but paired comparisons showed separation. Relative to PT, ED improved ARI by +0.028 (*p* = 2 × 10^−6^), VQT by +0.011 (*p* = 0.014), and VarQITE by +0.010 (*p* = 0.036); greedy VB and the annealing-only control were slightly worse, with intervals spanning zero (Fig. 5d). The median ARI rose from 0.46 to 0.49, consistent with the diagnostic’s predicted margin.

**Figure 5:**
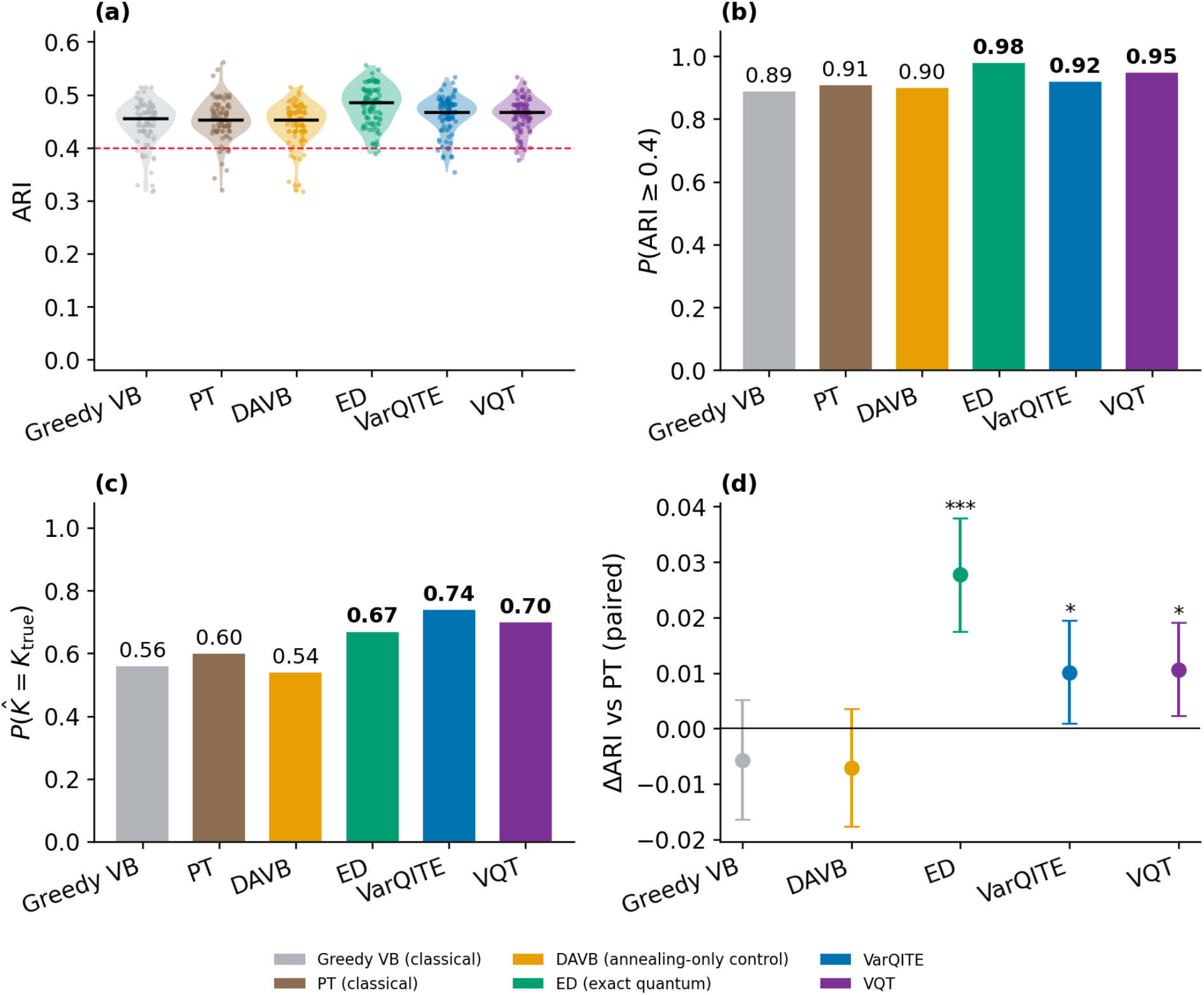
The quantum-coupled approach is more reliable than classical inference on real soil-microbiome data. Benchmark on the 88-soils continental gradient, clustering 89 samples by their pH into *K*_true_ = 4 balanced quantile groups over the 199 OTUs with *R* = 100 paired random seeds. (a) Per-seed adjusted Rand index (ARI); points are individual seeds, the horizontal bar is the median, and the dashed line marks the success threshold 0.4. (b) Single-run success probability *P* (ARI ≥ *ϵ*|*ϵ* = 0.4). (c) Success probability of estimating the number of clusters *P* 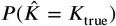. (d) Paired effect size relative to PT (ΔARI = ARI_method_ − ARI_PT_, matched by seed); markers show the mean, error bars the 95% bootstrap confidence interval, and annotations the paired Wilcoxon signed-rank significance. All three quantum methods significantly exceed PT (ED +0.028, *p* = 2 × 10^−6^(***); VQT +0.011, *p* = 0.014(*); VarQITE +0.010, *p* = 0.036(*)).

Assessing recovery of the cluster count distinguished the methods more distinctly, as in the synthetic setting. The quantum methods recovered *K*_true_ = 4 for 0.67 to 0.74 of seeds, compared with 0.54 to 0.60 for the classical baselines, and the strongest quantum method (VarQITE, 0.74 with Wilson 95% [0.65, 0.82]) did not overlap with the weakest classical method (DAVB, 0.54 with Wilson 95% [0.44, 0.63]). The relative ranking within the quantum methods mirrors a pattern observed twice on synthetic data: at a fixed operating point, the two circuit-based state preparations outperformed the exact preparation on this metric, while the opposite occurs when hyperparameters are sampled at random (Supplementary Material Table S1).

All six feature-selection methods gave identical results (Fig. 6): per-OTU selection distributions matched, and the top 20 discriminative OTUs were the same taxa in the same order. This is expected because quantum coupling affects only the assignment step, not the selection update, so we report it as a check rather than a result. Biologically, the selected set is dominated by Acidobacteria subgroups plus *Bradyrhizobium* and *Rubrobacter*, reproducing the pH-response pattern in the source study [38]. The model thus recovered coherent biological structure from untuned data.

**Figure 6:**
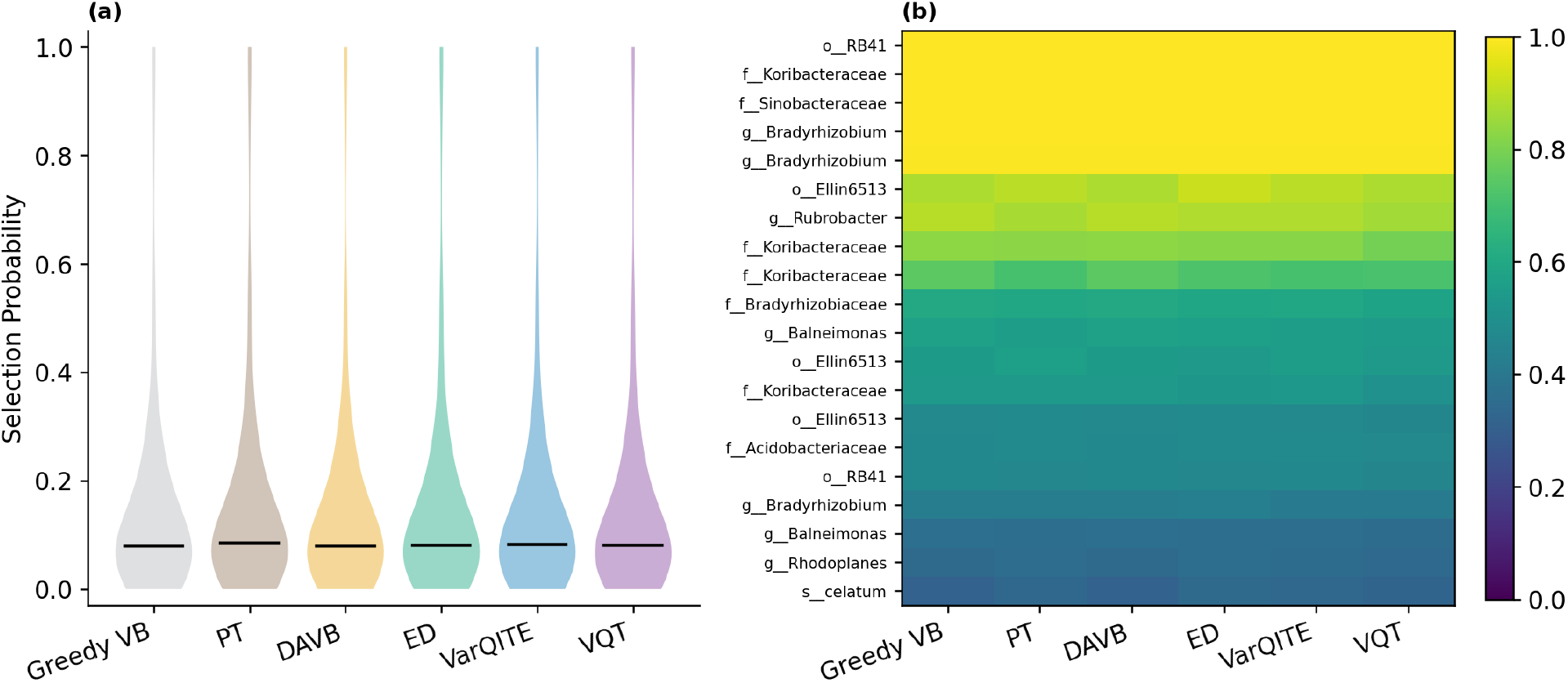
All methods recover the same biologically coherent pH-indicator taxa. Per-OTU selection probability from the DMM-SVVS selection variable, computed on the 88-soils pH data (89 samples, 199 OTUs, *K*_true_ = 4). (a) Distribution of per-OTU selection strength for each method; the horizontal bar is the median. (b) The top-20 discriminative OTUs, ranked by the ED approach and shown for every method.

**Figure 7:**
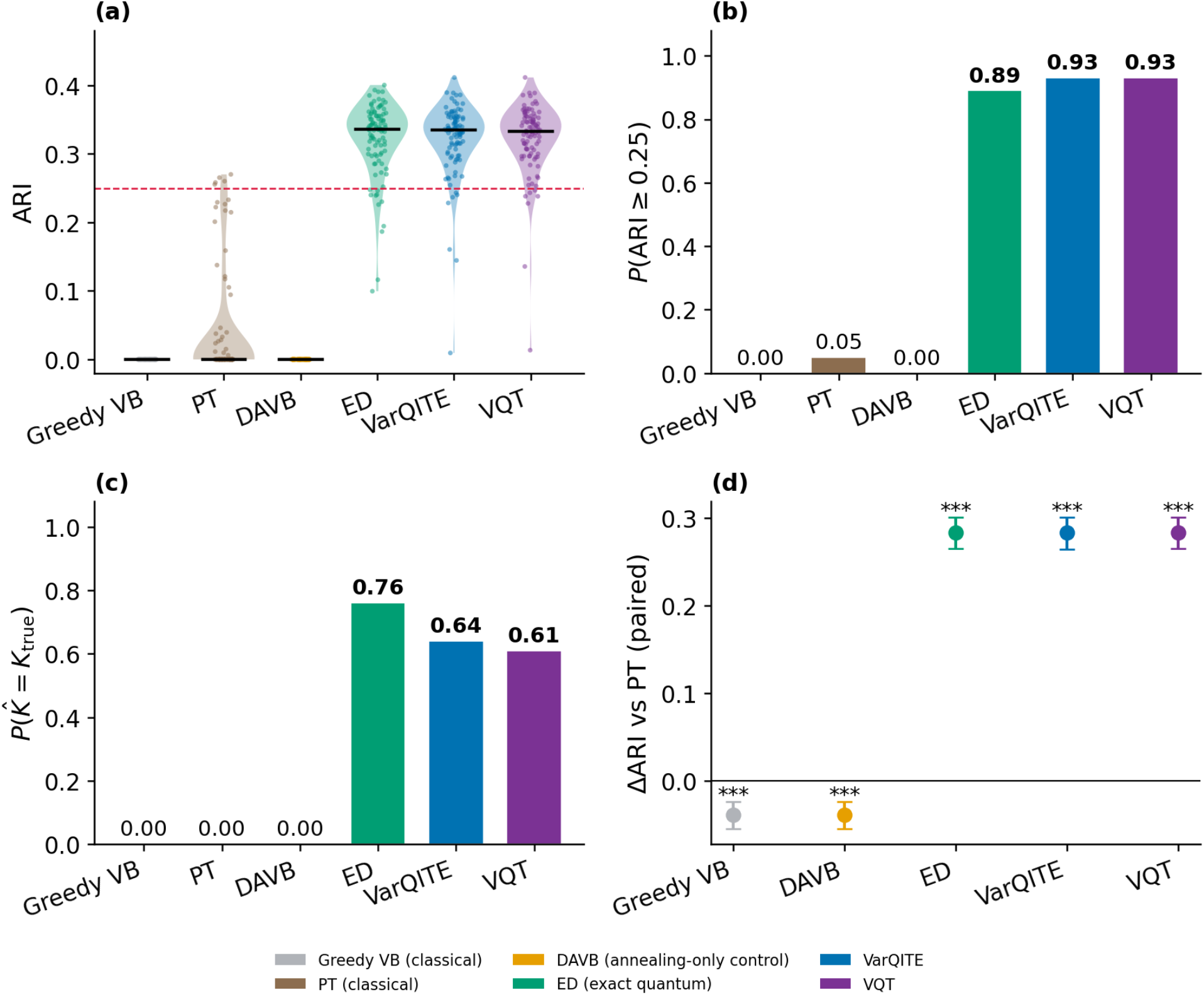
Human oral microbiome compositions controlled to the advantage regime. A human oral-microbiome dataset (HMP subgingival plaque, supragingival plaque, and tongue dorsum; *N* = 1149, *K* = 3) is placed in the advantage band by diluting its between-subsite signal to *σ*_min_ = 0.30. (a) Per-seed adjusted Rand index (ARI); points are individual seeds, the horizontal bar is the median, and the dashed line marks the success threshold 0.25. (b) Single-run success probability *P* (ARI ≥ *ϵ*|*ϵ* = 0.25). (c) Success probability of estimating the number of clusters 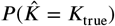. (d) aired ΔARI relative to the strong classical PT baseline with 95% bootstrap confidence intervals: all three quantum arms exceed PT by +0.284 (*p* ≈ 4× 10^−18^, Wilcoxon signed-rank).

#### 4.2.2. A controlled human cohort inside the advantage regime

The cohorts examined above lie outside the regime in which we predict a substantial effect, and this is not an accident of selection. Screening the human datasets available to us with the diagnostic of Section 3.1, body-site labels give *σ*_min_ between 0.85 and 0.99 [39, 46, 47], deep in the signal-dominated regime, while the disease labels we examined give *σ*_min_ ≈ 0 [48, 49, 50], below any detectability floor. This finding pertains to the datasets rather than the method, and it constrains the current usefulness of the method.

To evaluate our predictions on genuine human-derived compositions, we created a controlled dilution setting. Using HMP oral samples from three clusters (*N* = 1149, *K* = 3, *S* = 150) [39], we replaced each sample profile with a convex combination of the original profile and the globally pooled profile, 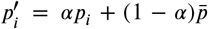, and then re-sampled to match its original library size. The mixing parameter *α* ranges from the unmodified data (*α* = 1) to a state where the between-class signal is fully eliminated (*α* = 0); we tuned *α* via bisection to achieve a desired signal fraction. Achieving *σ*_min_ ≈ 0.30 required *α* = 0.061, meaning about 94% of each profile was replaced by the pooled mean and the between-subsite signal was largely suppressed. The taxa, library sizes, sparsity patterns, and within-subsite sampling variability remained realistic, but the class separation did not.

At this difficulty, methods separate on endpoint-based metrics (no threshold). Over *R* = 100 paired seeds, mean ARI was 0.000 for greedy VB, 0.039 for PT, and 0.000 for the annealing-only control, versus 0.322 for all three quantum approaches. Cluster-count recovery was clearer: classical methods always collapsed to 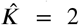 and never found the third cluster (0.00, Wilson 95% [0.00, 0.04]), while quantum approaches recovered 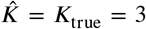 in 61–76 of 100 runs (fixed parameters: *K*_max_ = 4, DP concentration = 0.03, selection prior = 0.55, prune threshold = 0.02, *τ*_1_ = 86, *τ*_2_ = 173), with intervals disjoint from the classical bound. With parallel tempering, each quantum approach gained +0.284 ARI, with bootstrap intervals excluding zero. Because clusters overlap, achievable ARI is capped near 0.41, soa 0.40 threshold would mark all methods as failures; at τ = 0.25, success is 0.00–0.05 for classical methods versus 0.89–0.93 for quantum approaches. The annealing-only control behaved as expected. DAVB matched greedy VB exactly (0.000 mean ARI; Δ = −0.039 vs PT), indicating the separation is not due to the temperature schedule.

## 5. Conclusions and outlook

Where the signal fraction places a dataset in the advantage band, the separation between QBayMic and classical inference is categorical rather than incremental. In the benchmark cell, greedy VB, DAVB, and PT recovered the correct structure in none of the 100 runs, whereas QBayMic recovered it in 47 to 64; beyond three groups, the classical methods ceased to recover the number of clusters; and on a human-derived cohort placed in the band by controlled dilution, they succeeded in 0 to 5% of runs against 89 to 93%. These results support the reliability of the study. The accuracy ceiling remains unchanged; every method reaches it at high signal fractions, and the difference lies in the frequency of a single run.

Classical and quantum updates use the same per-assignmen energies; the difference is that the quantum energy operator includes off-diagonal terms. A softmax, at any temperature, only reweights the *K* assignments independently, so it can sharpen or smooth probabilities but cannot move mass between assignments. The transverse field instead couples assignments via single-qubit flips on a register whose connectivity does not weaken with size: assignments differing by one bit are directly linked, and any two of the *K* differ by at most ⌈log *K*⌉ bits. Scaling the energy operator by (1 − *s*) removes, rather than resolves, competition: early in the schedule the assignment energies are absent, so the softening never has to overcome energy differences of order 10^4^ nats with an inverse temperature of order 10. With the mixing term set to zero, the method reduces exactly to the classical update, making standard variational inference a limiting case; the question is whether intermediate schedules help. They do, over a range predicted in advance by the barrier decomposition, so the gain is not merely empirical.

The pattern of connectivity is a configurable design choice, not a fixed property of the method, so quantum circuit geometry becomes a substantive variable to study rather than a mere implementation detail. Our evidence that it has an effect is still preliminary, yet it is consistent. A single fixed circuit, without tuning to any specific cell, delivered the strong recovery of cluster counts across all conditions we evaluated, beating every fixed setting of an entropy-floor baseline; moreover, the best entropy-floor level depends on the cell, and bad choices can be expensive, reducing one cell’s score from 0.93 to 0.20. Direct measurements also eliminate the simplest explanation: the circuit’s E-step entropy aligns with that of the exact method, indicating the circuit is not merely functioning as a softening mechanism.

The standard remedies break down for different reasons. Restarts do not fail in the search itself but at the selection step: in this regime, free energy is negatively correlated with accuracy, so picking restarts by free energy consistently favors the wrong candidates. Deterministic annealing reaches the same partition as greedy ascent in every cell we tested; its operator stays diagonal at all temperatures, meaning the schedule may change the trajectory but not the final solution, and both approaches succeed or fail together throughout the sweep. Parallel tempering collapses even sooner, because its replicas never diverge enough to meaningfully exchange information.

One aspect of the construction warrants explicit discussion. Because the transverse-field term couples computational basis states that differ by a single bit, the effective coupling between any two candidate clusters depends on the specific binary encoding used to label those clusters. In the present representation, the *K*_max_ candidate clusters are encoded on ⌈log *K* ⌉_max_ qubits. For the reference configuration with *K*_max_ = 4 (*K*_true_ = 3), all four candidate clusters are mapped to the basis states |00⟩, |01⟩, |10⟩, |11⟩, and the procedure prunes to 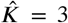. Under this encoding, pairs of states that differ by one bit (e.g., |00⟩ – |01⟩ and |00⟩ – |10⟩) are directly coupled, whereas pairs that differ by two bits (e.g., |00⟩ – |11⟩ and |01⟩ – |10⟩) interact only via second-order processes. Since this bit assignment is arbitrary, the resulting dynamics are not invariant under relabelling. We quantified the magnitude of this label-dependence by repeating the reference configuration over 15 random permutations of the cluster labels. This yielded a standard deviation of 0.029 in single-run reliability (*P* (ARI > 0.4) and 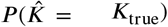 in Supplementary Material Table S7). This variability is small, approximately 4% of the 0.73 performance separation relative to the classical methods, and therefore does not affect the overall ranking of methods. Notably, the same encoding-induced structure implies that the mixer choice constitutes a genuine design parameter rather than an implementation detail; accordingly, Section 4.1.3 presents a preliminary comparison of two mixer geometries.

However, several limitations of this study should be considered. No natural cohort screened fell inside the advantage band; therefore, the human-derived dataset was placed there by dilution and is semi-synthetic; we do not claim that natural datasets in this regime are common. The benefit is not confined to the band; however, since on the soil dataset, which is signal-dominated, the quantum approaches still recovered the cluster count in 67 to 74% of runs against 54 to 60%. All results are exact noiseless simulations of a small number of qubit registers, which are classically tractable; therefore, we claim no speed-up. The deployable circuit preparation attained approximately three-quarters of the reliability of the exact preparation. Moreover, for unlabelled data, the diagnostic requires a preliminary partition, which is least dependable in the low-signal regime, where it matters most.

The future directions are as follows, each with an established route. Executing the assignment step on hardware at a mixer-active point of the schedule would test whether the advantage survives device noise; quantum-enhanced Markov chain Monte Carlo has already been run on superconducting hardware for a related sampling task [51, 52, 53], and zero-noise extrapolation offers a mitigation route for the readout [54, 55, 56]. A Hamiltonian-structured ansatz may close the preparation gap, because our annealed operator already has the alternating problem-and-mixer form that such ansatze exploit [57, 58], whereas the ansatz used here is generic. A wider screen over curated cohorts [59, 60] would establish how common the advantage band is in practice, which determines the reach of the method. Extending the quantum coupling to the feature-selection layer would substantially enlarge the register, raising trainability questions that do not arise at the sizes used here [61, 62, 63].

## Funding

This work was partly supported by JSPS KAKENHI Grant Number JP20H03240, JSPS KAKENHI Grant Number JP24K15175, JSPS KAKENHI Grant Number JP25K02261 and JST CREST Grant Number JPMJCR2231, Japan.

## Declaration of competing interest

The authors declare that they have no known competing financial interests or personal relationships that could have appeared to influence the work reported in this paper.

## Declaration of generative AI and AI-assisted technologies in the writing process

During the preparation of this work the authors used Claude Pro to improve the readability and language of the manuscript. After using this tool, the authors reviewed and edited the content as needed and take full responsibility for the content of the publication.

## Data availability

The source code and processed datasets used in this study are publicly available at https://github.com/tungtokyo1108/QBayMic

## CRediT authorship contribution statement

**Tung Dang:** Conceptualization, Methodology, Software, Validation, Formal analysis, Visualization, Writing – original draft, Writing – review & editing. **Artem Lysenko:** Conceptualization, Methodology, Investigation, Supervision, Writing – review & editing. **Tatsuhiko Tsunoda:** Conceptualization, Methodology, Investigation, Supervision, Funding acquisition, Project administration, Writing – review & editing.

## Notes

### Competing Interest Statement

The authors have declared no competing interest.

## References

[1] Rob Knight, Alison Vrbanac, Bryn C Taylor, Alexander Aksenov, Chris Callewaert, Justine Debelius, Antonio Gonzalez, Tomasz Kosciolek, Laura-Isobel McCall, Daniel McDonald, et al. Best practices for analysing microbiomes. Nature reviews microbiology, 16(7):410–422, 2018.

[2] Nicola Segata, Jacques Izard, Levi Waldron, Dirk Gevers, Larisa Miropolsky, Wendy S Garrett, and Curtis Huttenhower. Metagenomic biomarker discovery and explanation. Genome biology, 12(6):R60, 2011.

[3] Kris Sankaran and Susan P Holmes. Latent variable modeling for the microbiome. Biostatistics, 20(4):599–614, 2019.

[4] Ian Holmes, Keith Harris, and Christopher Quince. Dirichlet multinomial mixtures: generative models for microbial metagenomics. PloS one, 7(2):e30126, 2012.

[5] Yushu Shi, Liangliang Zhang, Christine B Peterson, Kim-Anh Do, and Robert R Jenq. Performance determinants of unsupervised clustering methods for microbiome data. Microbiome, 10(1):25, 2022.

[6] Tung Dang, Kie Kumaishi, Erika Usui, Shungo Kobori, Takumi Sato, Yusuke Toda, Yuji Yamasaki, Hisashi Tsujimoto, Yasunori Ichihashi, and Hiroyoshi Iwata. Stochastic variational variable selection for high-dimensional microbiome data. Microbiome, 10(1):236, 2022.

[7] Frederick A Matsen IV. Phylogenetics and the human microbiome.Systematic Biology, 64(1):e26–e41, 2015.

[8] Mengyu He, Ni Zhao, and Glen A Satten. Midasim: a fast and simple simulator for realistic microbiome data. Microbiome, 12(1):135, 2024.

[9] Matthew D Hoffman, David M Blei, Chong Wang, and John Paisley. Stochastic variational inference. Journal of machine learning research, 2013.

[10] David M Blei, Alp Kucukelbir, and Jon D McAuliffe. Variational inference: A review for statisticians. Journal of the American statistical Association, 112(518):859–877, 2017.

[11] Cheng Zhang, Judith Bütepage, Hedvig Kjellström, and Stephan Mandt. Advances in variational inference. IEEE transactions on pattern analysis and machine intelligence, 41(8):2008–2026, 2018.

[12] Ziang Chen, Yingzhou Li, and Jianfeng Lu. On the global convergence of randomized coordinate gradient descent for nonconvex optimization. SIAM Journal on Optimization, 33(2):713–738, 2023.

[13] Shunta Akiyama. Block coordinate descent for neural networks provably finds global minima. Advances in Neural Information Processing Systems, 38:158269–158308, 2026.

[14] Christophe Biernacki, Gilles Celeux, and Gérard Govaert. Choosing starting values for the em algorithm for getting the highest likelihood in multivariate gaussian mixture models. Computational Statistics & Data Analysis, 41(3-4):561–575, 2003.

[15] Wojciech Kwedlo. A new random approach for initialization of the multiple restart em algorithm for gaussian model-based clustering. Pattern Analysis and Applications, 18(4):757–770, 2015.

[16] Logan Mathesen, Giulia Pedrielli, Szu Hui Ng, and Zelda B Zabin-sky. Stochastic optimization with adaptive restart: A framework for integrated local and global learning. Journal of Global Optimization, 79(1):87–110, 2021.

[17] Kenneth Rose. Deterministic annealing for clustering, compression, classification, regression, and related optimization problems. Proceedings of the IEEE, 86(11):2210–2239, 1998.

[18] Kentaro Katahira, Kazuho Watanabe, and Masato Okada. Deterministic annealing variant of variational bayes method. In Journal of Physics: Conference Series, volume 95, page 012015, 2008.

[19] Chin-Wei Huang, Shawn Tan, Alexandre Lacoste, and Aaron Courville. Improving explorability in variational inference with annealed variational objectives. Advances in neural information processing systems, 31, 2018.

[20] David J Earl and Michael W Deem. Parallel tempering: Theory, applications, and new perspectives. Physical Chemistry Chemical Physics, 7(23):3910–3916, 2005.

[21] Malcolm Sambridge. A parallel tempering algorithm for probabilistic sampling and multimodal optimization. Geophysical Journal International, 196(1):357–374, 2014.

[22] Sayaka Shiota, Kei Hashimoto, Yoshihiko Nankaku, and Yi-Jian Wu. Deterministic annealing based training algorithm for bayesian speech recognition. In INTERSPEECH 2009 10th Annual Conference of the International Speech Communication Association, pages 680–683. Nagoya Institute of Technology, 2009.

[23] Anup Das. Deterministic and bayesian sparse signal processing algorithms for coherent multipath directions-of-arrival (doas) estimation. IEEE Journal of Oceanic Engineering, 44(4):1150–1164, 2018.

[24] Subinoy Adhikari and Jagannath Mondal. Elucidating protein dynamics through the optimal annealing of variational autoencoders. Journal of Chemical Theory and Computation, 21(13):6367–6379, 2025.

[25] Emma Prevot, Rory Toogood, Filippo Pagani, and Paul DW Kirk. Annealed variational mixtures for disease subtyping and biomarker discovery. Statistical Applications in Genetics and Molecular Biology, 25(1):20250001, 2026.

[26] Ulrich HE Hansmann. Parallel tempering algorithm for conformational studies of biological molecules. Chemical Physics Letters, 281(1-3):140–150, 1997.

[27] Satoshi Morita and Hidetoshi Nishimori. Mathematical foundation of quantum annealing. Journal of Mathematical Physics, 49(12), 2008.

[28] Sergio Boixo, Troels F Rønnow, Sergei V Isakov, Zhihui Wang, David Wecker, Daniel A Lidar, John M Martinis, and Matthias Troyer. Evidence for quantum annealing with more than one hundred qubits. Nature physics, 10(3):218–224, 2014.

[29] Philipp Hauke, Helmut G Katzgraber, Wolfgang Lechner, Hidetoshi Nishimori, and William D Oliver. Perspectives of quantum annealing: Methods and implementations. Reports on Progress in Physics, 83(5):054401, 2020.

[30] Sheir Yarkoni, Elena Raponi, Thomas Bäck, and Sebastian Schmitt. Quantum annealing for industry applications: Introduction and review. Reports on Progress in Physics, 85(10):104001, 2022.

[31] Issei Sato, Kenichi Kurihara, Shu Tanaka, Hiroshi Nakagawa, and Seiji Miyashita. Quantum annealing for variational bayes inference. arXiv preprint arXiv:0905.3528, 2009.

[32] Hideyuki Miyahara and Yuki Sughiyama. Quantum extension of variational bayes inference. Physical Review A, 98(2):022330, 2018.

[33] Hideyuki Miyahara and Vwani Roychowdhury. Quantum advantage in variational bayes inference. Proceedings of the National Academy of Sciences, 120(31):e2212660120, 2023.

[34] Guillaume Verdon, Jacob Marks, Sasha Nanda, Stefan Leichenauer, and Jack Hidary. Quantum hamiltonian-based models and the variational quantum thermalizer algorithm. arXiv preprint arXiv:1910.02071, 2019.

[35] Xiao Yuan, Suguru Endo, Qi Zhao, Ying Li, and Simon C Benjamin. Theory of variational quantum simulation. Quantum, 3:191, 2019.

[36] Sam McArdle, Tyson Jones, Suguru Endo, Ying Li, Simon C Benjamin, and Xiao Yuan. Variational ansatz-based quantum simulation of imaginary time evolution. npj Quantum Information, 5(1):75, 2019.

[37] Noam Slonim and Naftali Tishby. Agglomerative information bottleneck. Advances in neural information processing systems, 12, 1999.

[38] Christian L Lauber, Micah Hamady, Rob Knight, and Noah Fierer. Pyrosequencing-based assessment of soil ph as a predictor of soil bacterial community structure at the continental scale. Applied and environmental microbiology, 75(15):5111–5120, 2009.

[39] Structure, function and diversity of the healthy human microbiome.nature, 486(7402):207–214, 2012.

[40] Tadashi Kadowaki and Hidetoshi Nishimori. Quantum annealing in the transverse ising model. Physical Review E, 58(5):5355, 1998.

[41] Yuki Susa, Yu Yamashiro, Masayuki Yamamoto, and Hidetoshi Nishimori. Exponential speedup of quantum annealing by inhomogeneous driving of the transverse field. Journal of the Physical Society of Japan, 87(2):023002, 2018.

[42] Marco Cerezo, Andrew Arrasmith, Ryan Babbush, Simon C Benjamin, Suguru Endo, Keisuke Fujii, Jarrod R McClean, Kosuke Mitarai, Xiao Yuan, Lukasz Cincio, et al. Variational quantum algorithms. Nature Reviews Physics, 3(9):625–644, 2021.

[43] Ville Bergholm et al. Pennylane: Automatic differentiation of quantum circuits. arXiv preprint arXiv: 1811.04968, 2018.

[44] Jianhua Lin. Divergence measures based on the shannon entropy.IEEE Transactions on Information theory, 37(1):145–151, 1991.

[45] Kenneth Rose, Eitan Gurewitz, and Geoffrey Fox. A deterministic annealing approach to clustering. Pattern Recognition Letters, 11(9):589–594, 1990.

[46] A framework for human microbiome research. nature, 486(7402):215–221, 2012.

[47] Jun Hang, Valmik Desai, Nela Zavaljevski, Yu Yang, Xiaoxu Lin, Ravi Vijaya Satya, Luis J Martinez, Jason M Blaylock, Richard G Jarman, Stephen J Thomas, et al. 16s rrna gene pyrosequencing of reference and clinical samples and investigation of the temperature stability of microbiome profiles. Microbiome, 2(1):31, 2014.

[48] Catherine A Lozupone, Marcella Li, Thomas B Campbell, Sonia C Flores, Derek Linderman, Matthew J Gebert, Rob Knight, Andrew P Fontenot, and Brent E Palmer. Alterations in the gut microbiota associated with hiv-1 infection. Cell host & microbe, 14(3):329–339, 2013.

[49] Julia K Goodrich, Jillian L Waters, Angela C Poole, Jessica L Sutter, Omry Koren, Ran Blekhman, Michelle Beaumont, William Van Treuren, Rob Knight, Jordana T Bell, et al. Human genetics shape the gut microbiome. Cell, 159(4):789–799, 2014.

[50] Georg Zeller, Julien Tap, Anita Y Voigt, Shinichi Sunagawa, Jens Roat Kultima, Paul I Costea, Aurélien Amiot, Jürgen Böhm, Francesco Brunetti, Nina Habermann, et al. Potential of fecal microbiota for early-stage detection of colorectal cancer. Molecular systems biology, 10(11):MSB145645, 2014.

[51] David Layden, Guglielmo Mazzola, Ryan V Mishmash, Mario Motta, Pawel Wocjan, Jin-Sung Kim, and Sarah Sheldon. Quantum-enhanced markov chain monte carlo. Nature, 619(7969):282–287, 2023.

[52] Alev Orfi and Dries Sels. Bounding the speedup of the quantum-enhanced markov-chain monte carlo algorithm. Physical Review A, 110(5):052414, 2024.

[53] Stuart Ferguson and Petros Wallden. Quantum-enhanced markov chain monte carlo for systems larger than a quantum computer. Physical Review Research, 7(1):013231, 2025.

[54] Kristan Temme, Sergey Bravyi, and Jay M Gambetta. Error mitigation for short-depth quantum circuits. Physical review letters, 119(18):180509, 2017.

[55] Martin Larocca, Supanut Thanasilp, Samson Wang, Kunal Sharma, Jacob Biamonte, Patrick J Coles, Lukasz Cincio, Jarrod R McClean, Zoë Holmes, and Marco Cerezo. Barren plateaus in variational quantum computing. Nature Reviews Physics, 7(4):174–189, 2025.

[56] Amira Abbas, Andris Ambainis, Brandon Augustino, Andreas Bärtschi, Harry Buhrman, Carleton Coffrin, Giorgio Cortiana, Vedran Dunjko, Daniel J Egger, Bruce G Elmegreen, et al. Challenges and opportunities in quantum optimization. Nature Reviews Physics, 6(12):718–735, 2024.

[57] Edward Farhi, Jeffrey Goldstone, and Sam Gutmann. A quantum approximate optimization algorithm. arXiv preprint arXiv:1411.4028, 2014.

[58] Dave Wecker, Matthew B Hastings, and Matthias Troyer. Progress towards practical quantum variational algorithms. Physical Review A, 92(4):042303, 2015.

[59] Edoardo Pasolli, Lucas Schiffer, Paolo Manghi, Audrey Renson, Valerie Obenchain, Duy Tin Truong, Francesco Beghini, Faizan Malik, Marcel Ramos, Jennifer B Dowd, et al. Accessible, curated metagenomic data through experimenthub. Nature methods, 14(11):1023–1024, 2017.

[60] Claire Duvallet, Sean M Gibbons, Thomas Gurry, Rafael A Irizarry, and Eric J Alm. Meta-analysis of gut microbiome studies identifies disease-specific and shared responses. Nature communications, 8(1):1784, 2017.

[61] Jarrod R McClean, Sergio Boixo, Vadim N Smelyanskiy, Ryan Babbush, and Hartmut Neven. Barren plateaus in quantum neural network training landscapes. Nature communications, 9(1):4812, 2018.

[62] Amira Abbas, David Sutter, Christa Zoufal, Aurélien Lucchi, Alessio Figalli, and Stefan Woerner. The power of quantum neural networks. Nature computational science, 1(6):403–409, 2021.

[63] Martin Larocca, Nathan Ju, Diego García-Martín, Patrick J Coles, and Marco Cerezo. Theory of overparametrization in quantum neural networks. Nature Computational Science, 3(6):542–551, 2023.

