## Supplementary materials for "QBayMic: Quantum-coupled variational Bayes for clustering and feature selection in low-signal microbiome data"

### Appendix A Supplementary Tables and Figures

**Table S1: Hyperparameter robustness of the quantum-coupled E-step at the reference cell.** Distribution of the adjusted Rand index (ARI) and of the recovered cluster count over 100 random hyperparameter draws per method (shared master seed) at the low-signal reference cell ($\sigma \approx 0.29$; 400 samples $\times$ 5000 OTUs, separation 0.2, zero-inflation 0.8; over-specified truncation $K_{\mathrm{max}} \in \{4, 5, 6\} \geq K_{\mathrm{true}} = 3$). Each draw is an independent hyperparameter sample, so no label information enters any single run. This sensitivity analysis complements Fig. 2, which reports single-run reliability at a fixed operating point selected label-free; absolute values are lower here because the random search includes deliberately poor configurations. Even averaged over the hyperparameter prior, the quantum E-steps raise single-run success $P(\mathrm{ARI} > 0.4)$ from 0.02 to 0.23–0.42 and true-$K$ recovery from 0.01 to 0.20–0.35, so the advantage is not an artifact of one tuned configuration.

| Method | median | $(Q_1, Q_3)$ | mean | max | $P(\mathrm{ARI}{>}0.3)$ | $P(\mathrm{ARI}{>}0.4)$ | $P(\hat{K}{=}3)$ |
|---|---|---|---|---|---|---|---|
| Greedy VB | −0.00 | (−0.00, 0.00) | 0.02 | 0.44 | 0.04 | 0.02 (0.01–0.07) | 0.01 |
| QBayMic (ED, exact) | 0.38 | (0.22, 0.44) | 0.35 | 0.66 | 0.65 | 0.42 (0.33–0.52) | 0.35 |
| QBayMic (VarQITE) | 0.24 | (0.13, 0.36) | 0.27 | 0.67 | 0.37 | 0.23 (0.16–0.32) | 0.24 |
| QBayMic (VQT) | 0.24 | (0.14, 0.40) | 0.28 | 0.68 | 0.41 | 0.25 (0.18–0.34) | 0.20 |

Parentheses after $P(\mathrm{ARI}{>}0.4)$ are Wilson 95% confidence intervals ($n = 100$). $\hat{K}$ is the recovered occupied-component count under a fixed, method-agnostic occupancy rule; $K_{\mathrm{true}} = 3$. At the fixed operating point of Fig. 2c the scalable methods recover $K$ best; averaged over random hyperparameters (above) the exact E-step is most robust.

**Table S2: Pairwise comparisons at the reference cell.** Effect sizes for the random-hyperparameter distributions in Table S1. All three quantum E-steps dominate greedy VB by a large margin. Among the quantum variants the exact E-step (ED) modestly but significantly exceeds both scalable preparations, quantifying the cost of approximate Gibbs preparation, while VQT and VarQITE are statistically indistinguishable (confirming they are independent methods that converge to the same Gibbs state). Effect sizes are the primary evidence; $p$-values are secondary.

| Comparison | $\Delta$median ARI | 95% CI | test | $p$ |
|---|---|---|---|---|
| ED − VB | +0.38 | n/a | Mann–Whitney | $5 \times 10^{-31}$ |
| VarQITE − VB | +0.24 | n/a | Mann–Whitney | $1 \times 10^{-26}$ |
| VQT − VB | +0.24 | n/a | Mann–Whitney | $8 \times 10^{-26}$ |
| ED − VarQITE | +0.07 | [+0.03, +0.10] | paired Wilcoxon | $1 \times 10^{-5}$ |
| ED − VQT | +0.07 | [+0.03, +0.11] | paired Wilcoxon | $3 \times 10^{-5}$ |
| VQT − VarQITE | +0.00 | [ 0.00, 0.00] | paired Wilcoxon | 0.28 |

ED, VarQITE and VQT share identical hyperparameter and seed streams, so their comparisons are paired (Wilcoxon) with a bootstrap 95% CI on the median difference. Greedy VB uses an independent stream (it draws fewer parameters per trial); VB comparisons are unpaired (Mann–Whitney) and a paired CI is not defined (n/a).

**Table S3: Metric robustness: the same ranking holds under normalized mutual information (NMI).** Distribution of NMI over the same 100 random hyperparameter draws as Table S1, at the reference cell. The ordering and every pairwise conclusion match the ARI analysis: greedy VB sits near chance, the scalable E-steps cluster together, and the exact E-step (ED) leads, with ED exceeding both VarQITE and VQT (paired Wilcoxon, +0.05, $p \approx 2 \times 10^{-3}$) and VQT indistinguishable from VarQITE (+0.00, $p = 0.69$).

| Method | median | ($Q_1$, $Q_3$) | mean | max | $P$(NMI>0.3) | $P$(NMI>0.4) |
|---|---|---|---|---|---|---|
| Greedy VB | 0.00 | (0.00, 0.02) | 0.03 | 0.36 | 0.03 | 0.00 |
| QBayMic (ED, exact) | 0.37 | (0.23, 0.48) | 0.35 | 0.62 | 0.64 | 0.48 |
| QBayMic (VarQITE) | 0.32 | (0.18, 0.43) | 0.31 | 0.66 | 0.53 | 0.30 |
| QBayMic (VQT) | 0.32 | (0.17, 0.43) | 0.31 | 0.66 | 0.53 | 0.35 |

**Table S4: Hyperparameter search specification.** Ranges indicate parameters searched (100 random draws, shared master seed) on the low-signal reference cell ($\sigma \approx 0.29$; 400 samples × 5000 OTUs, separation 0.2, zero-inflation 0.8; over-specified truncation $K_{\text{true}} = 3$). "=" indicates a fixed value; "n/a" indicates a method without that parameter. Shared model parameters use the same ranges across methods. Annealing-schedule parameters use the same ranges across the three annealed methods and have no greedy-VB analog. Quantum-architecture parameters were fixed at standard defaults and not tuned on this cell. The exact method (ED) has no architecture parameters, so its advantage is independent of architecture tuning.

| Hyperparameter | Greedy VB | ED (exact) | VarQITE | VQT |
|---|---|---|---|---|
| *Shared model parameters (searched, identical ranges)* | | | | |
| $K_{\max}$ | $[4, 6]$ | $[4, 6]$ | $[4, 6]$ | $[4, 6]$ |
| DP concentration | $[0.1, 10]$ | $[0.1, 10]$ | $[0.1, 10]$ | $[0.1, 10]$ |
| selection prior | $[0.1, 0.95]$ | $[0.1, 0.95]$ | $[0.1, 0.95]$ | $[0.1, 0.95]$ |
| prune threshold | $[0.05, 0.5]$ | $[0.05, 0.5]$ | $[0.05, 0.5]$ | $[0.05, 0.5]$ |
| *Annealing schedule (searched, identical across annealing methods)* | | | | |
| quantum-phase length $\tau_1$ | n/a | $[40, 110]$ | $[40, 110]$ | $[40, 110]$ |
| convergence guard offset $\tau_2$ ($\delta$) | n/a | $[90, 220]$ | $[90, 220]$ | $[90, 220]$ |
| initial inverse temperature $\beta_0$ | n/a | $= 30$ | $= 30$ | $= 30$ |
| initial mixer weight $s_0$ | n/a | $= 1$ | $= 1$ | $= 1$ |
| mixer strength $\Gamma$ | n/a | $1 \to 0$ | $1 \to 0$ | $1 \to 0$ |
| *Quantum architecture (fixed defaults, not tuned on the cell)* | | | | |
| ansatz depth | n/a | n/a | $= 3$ | $= 3$ |
| thermalization steps | n/a | n/a | $= 40$ | $= 40$ |
| learning rate | n/a | n/a | n/a$^{\ddagger}$ | $= 0.05$ |
| mixer | n/a | transverse | transverse | transverse |
| *Optimization budget* | | | | |
| max EM iterations | $= 400$ | $= 400$ | $= 400$ | $= 400$ |
| convergence tol. | $= 10^{-4}$ | $= 10^{-4}$ | $= 10^{-4}$ | $= 10^{-4}$ |

$^{\ddagger}$VarQITE uses imaginary-time evolution (step $d\tau = 0.2$) rather than a gradient learning rate. Other numerical settings (prune start/interval, trigamma correction) were fixed and identical across the quantum methods.

**Table S5: Sampling uncertainty for the regime sweep (Fig. 3).** For each cell (sorted by signal fraction $\sigma$) the single-run success rate $P(\text{ARI} > 0.4) = k/R$ is reported with its Wilson score 95% confidence interval $[\,\text{lo}, \text{hi}\,]$ from $R = 15$ paired seeds, for greedy VB, the annealing-only control (DAVB), parallel tempering (PT), the exact quantum E-step (QBayMic-ED), and the deployable variational quantum circuit (VQT). At $R = 15$ each interval is $\approx 0.2$–$0.5$ wide, so adjacent-cell differences within a method are generally not resolvable; in particular the apparent QBayMic dip from 0.73 ($\sigma = 0.291$) to 0.53 ($\sigma = 0.305$) has overlapping intervals $[0.48, 0.89]$ vs. $[0.30, 0.75]$ and is not a resolved feature. DAVB equals greedy VB in every cell (deterministic annealing does not change the reached basin on this landscape; see main text). VQT, the runnable circuit, tracks QBayMic-ED across all sixteen cells (its intervals overlap ED's in every cell), confirming the advantage regime carries over to a real ansatz: it wins where ED wins (advantage cells $\sigma = 0.291$ and 0.388–0.427) and saturates to 1.00 with ED in the signal-dominated regime ($\sigma \geq 0.56$). †: advantage cell (the quantum arm exceeds the best classical arm by $\geq 0.30$).

| | | | $P(\text{ARI} > 0.4)$ with Wilson 95% CI | | | | |
|---|---|---|---|---|---|---|---|
| sep | ZI | $\sigma$ | VB | DAVB | PT | QBayMic | VQT |
| 0.2 | 0.9 | 0.194 | 0.00 [0.00, 0.20] | 0.00 [0.00, 0.20] | 0.00 [0.00, 0.20] | 0.00 [0.00, 0.20] | 0.00 [0.00, 0.20] |
| 0.2 | 0.8 | 0.291† | 0.00 [0.00, 0.20] | 0.00 [0.00, 0.20] | 0.00 [0.00, 0.20] | **0.73** [0.48, 0.89] | **0.80** [0.55, 0.93] |
| 0.3 | 0.9 | 0.305† | 0.00 [0.00, 0.20] | 0.00 [0.00, 0.20] | 0.00 [0.00, 0.20] | **0.53** [0.30, 0.75] | 0.27 [0.11, 0.52] |
| 0.2 | 0.7 | 0.361† | 0.40 [0.20, 0.64] | 0.40 [0.20, 0.64] | 0.07 [0.01, 0.30] | **0.80** [0.55, 0.93] | 0.60 [0.36, 0.80] |
| 0.4 | 0.9 | 0.388† | 0.07 [0.01, 0.30] | 0.07 [0.01, 0.30] | 0.33 [0.15, 0.58] | **0.87** [0.62, 0.96] | **0.87** [0.62, 0.96] |
| 0.2 | 0.6 | 0.419 | 1.00 [0.80, 1.00] | 1.00 [0.80, 1.00] | 0.67 [0.42, 0.85] | 1.00 [0.80, 1.00] | 1.00 [0.80, 1.00] |
| 0.3 | 0.8 | 0.427† | 0.27 [0.11, 0.52] | 0.27 [0.11, 0.52] | 0.40 [0.20, 0.64] | **0.93** [0.70, 0.99] | **0.93** [0.70, 0.99] |
| 0.5 | 0.9 | 0.476 | 0.47 [0.25, 0.70] | 0.47 [0.25, 0.70] | 0.73 [0.48, 0.89] | 1.00 [0.80, 1.00] | 0.93 [0.70, 0.99] |
| 0.3 | 0.7 | 0.509 | 1.00 [0.80, 1.00] | 1.00 [0.80, 1.00] | 0.93 [0.70, 0.99] | 1.00 [0.80, 1.00] | 1.00 [0.80, 1.00] |
| 0.4 | 0.8 | 0.528 | 0.87 [0.62, 0.96] | 0.87 [0.62, 0.96] | 0.80 [0.55, 0.93] | 1.00 [0.80, 1.00] | 1.00 [0.80, 1.00] |
| 0.3 | 0.6 | 0.562 | 1.00 [0.80, 1.00] | 1.00 [0.80, 1.00] | 1.00 [0.80, 1.00] | 1.00 [0.80, 1.00] | 1.00 [0.80, 1.00] |
| 0.5 | 0.8 | 0.610 | 1.00 [0.80, 1.00] | 1.00 [0.80, 1.00] | 1.00 [0.80, 1.00] | 1.00 [0.80, 1.00] | 1.00 [0.80, 1.00] |
| 0.4 | 0.7 | 0.618 | 1.00 [0.80, 1.00] | 1.00 [0.80, 1.00] | 0.93 [0.70, 0.99] | 1.00 [0.80, 1.00] | 1.00 [0.80, 1.00] |
| 0.4 | 0.6 | 0.674 | 1.00 [0.80, 1.00] | 1.00 [0.80, 1.00] | 1.00 [0.80, 1.00] | 1.00 [0.80, 1.00] | 1.00 [0.80, 1.00] |
| 0.5 | 0.7 | 0.696 | 1.00 [0.80, 1.00] | 1.00 [0.80, 1.00] | 1.00 [0.80, 1.00] | 1.00 [0.80, 1.00] | 1.00 [0.80, 1.00] |
| 0.5 | 0.6 | 0.750 | 1.00 [0.80, 1.00] | 1.00 [0.80, 1.00] | 1.00 [0.80, 1.00] | 1.00 [0.80, 1.00] | 1.00 [0.80, 1.00] |

**Table S6: Cluster-number recovery $P(\hat{K} = K_{\text{true}})$ across the exact E-step, a tuned entropy-regularization control, and quantum circuits ($K_{\text{true}} = 3$, R=15).** Two hard cells where the exact preparation over-clusters. ED: exact Gibbs E-step. softED($\eta^\star$): exact ED with a entropy floor $r \leftarrow (1-\eta)r + \eta/K$, reported at its *best* $\eta$ per cell (the value differs between cells, and a poor $\eta$ collapses recovery – e.g. $\eta = 0.5$ gives 0.20 at $\sigma$=0.29). $\text{VQT}_{\text{TF}}$: variational quantum circuit, transverse-field mixer (result is stable across ansatz depth 1–8 and 40–400 optimisation steps). $\text{VQT}_{\text{CS}}$: same circuit with a cyclic-shift mixer.

| | $\sigma = 0.43$ | $\sigma = 0.53$ |
|---|---|---|
| exact ED | 0.13 | 0.20 |
| softED ($\eta^\star$), best-tuned | 0.33 | 0.53 |
| $\text{VQT}_{\text{TF}}$ (fixed circuit) | 0.53 | 0.33 |
| $\text{VQT}_{\text{CS}}$ (mixer design) | 0.53 | **0.60** |

**Table S7: Reliability of the transverse-field VQT is invariant to the label-to-qubit map.** At the reference configuration ($\sigma = 0.291$, $K_{\text{true}} = 3$, $R = 15$ random initialisations per row; the model searches over $K_{\max} = 4$ candidate clusters carried on two qubits ($|00\rangle, |01\rangle, |10\rangle, |11\rangle$, no phantom padding) and prunes to $\hat{K} = 3$), single-run reliability $P(\text{ARI} > 0.4)$, cluster-number recovery $P(\hat{K} = K_{\text{true}})$, and mean ARI are reported for the identity label-to-basis-state map and for 15 random permutations of it. The permutation-induced spread (standard deviation across maps) is 0.029 in $P(\text{ARI} > 0.4)$ and 0.040 in $P(\hat{K})$ – about 4% of the $\approx 0.73$ gap to the classical arms at this cell – so the arbitrary assignment of clusters to computational basis states does not affect the ranking of methods. The same one-bit coupling structure of the transverse field is what makes the mixer a design variable.

| label map | $P(\text{ARI} > 0.4)$ | $P(\hat{K} = K_{\text{true}})$ | mean ARI |
|---|---|---|---|
| identity | 0.73 | 0.93 | 0.495 |
| perm 1 | 0.67 | 0.93 | 0.430 |
| perm 2 | 0.73 | 0.80 | 0.413 |
| perm 3 | 0.73 | 0.87 | 0.408 |
| perm 4 | 0.73 | 0.87 | 0.408 |
| perm 5 | 0.73 | 0.87 | 0.407 |
| perm 6 | 0.73 | 0.93 | 0.487 |
| perm 7 | 0.67 | 0.93 | 0.429 |
| perm 8 | 0.73 | 0.87 | 0.409 |
| perm 9 | 0.73 | 0.93 | 0.495 |
| perm 10 | 0.73 | 0.87 | 0.414 |
| perm 11 | 0.73 | 0.87 | 0.407 |
| perm 12 | 0.73 | 0.87 | 0.408 |
| perm 13 | 0.67 | 0.93 | 0.429 |
| perm 14 | 0.67 | 0.93 | 0.430 |
| perm 15 | 0.67 | 0.93 | 0.429 |
| mean ± SD (15 perms) | $0.71 \pm 0.03$ | $0.89 \pm 0.04$ | $0.427 \pm 0.028$ |
| range (15 perms) | $[0.67, 0.73]$ | $[0.80, 0.93]$ | $[0.41, 0.50]$ |

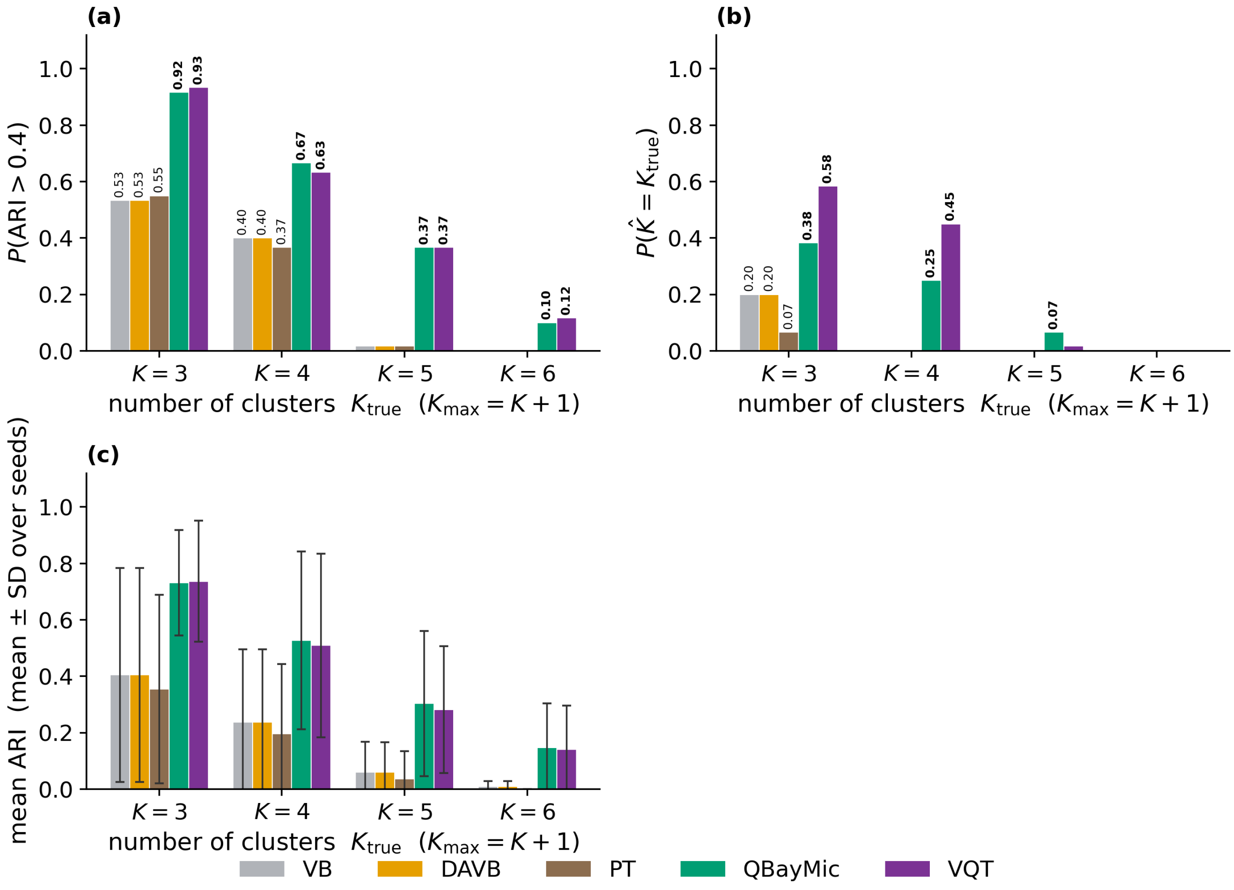

**Supplementary Figure S1**: The quantum-coupled E-step scales to more clusters, where classical inference collapses. Performance at increasing cluster number $K_{\mathrm{true}} \in \{3, 4, 5, 6\}$, cluster separation $\in \{0.2, 0.3, 0.4, 0.5\}$ with the truncation over-specified ($K_{\mathrm{max}} = K_{\mathrm{true}} + 1$, so the cluster count must be discovered). Each bar aggregates $R = 15$ paired seeds over four separation cells at matched compute (400 energy evaluations); only the E-step dynamics differ. (a) Single-run reliability $P(\mathrm{ARI} > 0.4)$ and (b) cluster-count discovery $P(\hat{K} = K_{\mathrm{true}})$: QBayMic leads at every $K$ and is the only method with non-zero cluster-count recovery beyond $K$=3, while all three classical methods fall to zero as $K$ grows. (c) Mean adjusted Rand index (mean ± SD over seeds).

### Appendix B  Generative model: full hierarchical and selection priors

We model the count matrix $\mathbf{X} \in \mathbb{Z}_{\geq 0}^{N\times S}$ with the Dirichlet–multinomial mixture with stochastic variational variable selection (DMM–SVVS) [1], one of the most widely used generative model for microbiome counts [2]. The main text keeps only the quantities a reader needs to follow the quantum E-step; the full model is as follows.

#### B.1  Mixture and stick-breaking prior

Each sample $i \in \{1, \dots, N\}$ carries a latent cluster label $z_i \in \{1, \dots, K_{\max}\}$ with $z_i \sim \text{Categorical}(\boldsymbol{\pi})$. The mixing weights follow a truncated stick-breaking (GEM) prior at truncation level $K_{\max}$,

$$v_k \sim \text{Beta}(1, \alpha_0), \qquad \pi_k = v_k \prod_{l<k}(1 - v_l), \qquad k = 1, \dots, K_{\max}, \tag{S1}$$

with $v_{K_{\max}} \equiv 1$ so that $\sum_k \pi_k = 1$. The stick-breaking prior shrinks unused components, so the number of occupied clusters is inferred rather than fixed; over-specifying $K_{\max} > K_{\text{true}}$ forces the model to discover the cluster count, the harder model-selection endpoint of the body.

#### B.2  Dirichlet–multinomial components

Conditional on $z_i = k$, the counts follow a Dirichlet–multinomial with cluster concentration $\boldsymbol{\phi}_k \in \mathbb{R}^S_{>0}$,

$$\mathbf{p}_i \mid z_i{=}k \sim \text{Dirichlet}(\boldsymbol{\phi}_k), \qquad \mathbf{x}_i \mid \mathbf{p}_i \sim \text{Multinomial}(n_i, \mathbf{p}_i), \qquad n_i = \textstyle\sum_s x_{is}, \tag{S2}$$

which after marginalising $\mathbf{p}_i$ is the Dirichlet–multinomial likelihood $p(\mathbf{x}_i \mid \boldsymbol{\phi}_k) = \binom{n_i}{\mathbf{x}_i} B(\mathbf{x}_i + \boldsymbol{\phi}_k)/B(\boldsymbol{\phi}_k)$, with $B(\cdot)$ the multivariate Beta function. The overdispersion of the Dirichlet–multinomial is what makes it the appropriate likelihood for microbiome counts [2].

#### B.3  Variable-selection layer

A per-taxon Bernoulli selection indicator $\gamma_s \in \{0, 1\}$, $\gamma_s \sim \text{Bernoulli}(\omega)$ with selection prior $\omega$, partitions taxa into an informative set ($\gamma_s{=}1$, cluster-specific concentration) and a shared background ($\gamma_s{=}0$, common concentration),

$$\phi_{ks} = \begin{cases} \phi_{ks}^{\text{sig}}, & \gamma_s = 1 \text{ (cluster-specific)}, \\ \phi_s^{\text{bg}}, & \gamma_s = 0 \text{ (shared)}. \end{cases} \tag{S3}$$

This layer makes the model appropriate for the high-dimensional, sparse regime, in which only a small fraction of taxa carry cluster signal. The conjugate hyperpriors on $(\boldsymbol{\phi}^{\text{sig}}, \boldsymbol{\phi}^{\text{bg}}, \omega, \alpha_0)$ follow the DMM–SVVS construction.

#### B.4  Mean-field family and the E-step we make quantum

Inference is over the joint posterior of $(\{z_i\}, \boldsymbol{\pi}, \{\boldsymbol{\phi}_k\}, \{\gamma_s\})$. The mean-field family factorises as [1, 3]

$$q = \Big[\textstyle\prod_i q(z_i)\Big]\, q(\boldsymbol{\pi})\, \Big[\textstyle\prod_k q(\boldsymbol{\phi}_k)\Big]\, \Big[\textstyle\prod_s q(\gamma_s)\Big], \tag{S4}$$

and coordinate ascent maximises the ELBO [1, 3]. The assignment factor $q(z_i)$ is the only update we replace; the remaining factors use the standard DMM–SVVS closed-form updates and are shared identically across all compared methods, so any measured difference is attributable to the E-step alone.

### Appendix C Three preparations of the quantum-coupled E-step

We realise $\rho_i(s)$ by three methods with an identical target. They fall into two categories that it is important to keep distinct. Exact diagonalization (ED) is a classical computation (a dense matrix exponential) that we use only as a ground-truth oracle; it does not run as a quantum algorithm and does not scale, because it manipulates the full $2^{n_q} \times 2^{n_q}$ density matrix explicitly. The variational quantum thermalizer (VQT) and variational imaginary-time evolution (VarQITE) are the genuine quantum preparations: each prepares $\rho_i(s)$ on a parameterised quantum circuit, manipulating only an $n_q$- (VQT) or $2n_q$-qubit (VarQITE) state vector and a small set of variational angles, with no explicit density matrix. This circuit-based preparation is the contribution that carries to near-term devices and that scales beyond the register sizes ED can reach: the quantum cost is set by circuit width, depth and parameter count (Tables S8–S9), not by the dimension of a stored matrix.

#### C.1 Exact preparation (exact diagonalization, ED)

ED yields the exact target thermal state and, in our implementation, prepares it on the same state-vector register the variational methods use, so that the whole pipeline is exercised end to end. It proceeds in two stages. *(i) Classical stage:* $\hat{H}_i(s)$ is formed as an explicit $2^{n_q} \times 2^{n_q}$ matrix and the thermal state $\rho_i(s) = e^{-\beta \hat{H}_i(s)}/Z_i$ is computed by dense matrix exponentiation and eigendecomposition, $\rho_i(s) = \sum_k \lambda_k \left|\phi_k\right\rangle\!\left\langle\phi_k\right|$. *(ii) State-vector stage:* the eigenpairs are assembled into a thermofield-double purification $|\Psi\rangle = \sum_k \sqrt{\lambda_k}\, |\phi_k\rangle_S \otimes |k\rangle_A$ on a $2n_q$-qubit system–ancilla register, loaded into a PennyLane state-vector simulator (`StatePrep`), and the reduced system state $\rho_S = \mathrm{Tr}_A |\Psi\rangle\langle\Psi|$ is recovered by partial trace; the responsibilities are its diagonal, $r_{ik} = \langle k|\, \rho_S \,|k\rangle$. This uses qubit registers and the same purification/readout as VarQITE, so the exact and variational preparations are compared on an identical footing.

The essential distinction from VQT and VarQITE is not the register but how the state is obtained: ED computes the Gibbs weights $\lambda_k$ classically and merely loads the resulting state, so it is a verification scaffold rather than a scalable quantum algorithm — no variational circuit generates the thermal state, and the classical diagonalization costs $\Theta(2^{2n_q})$ in memory and worse in time. ED is therefore tractable only at the small clustering register ($n_q = \lceil \log_2 K \rceil$) and does not scale, which is precisely why the variational, circuit-generated preparations are needed. Its role here is as the exact ground-truth oracle against which VQT and VarQITE are validated (Sec. Appendix D): when we say those methods “reproduce the exact E-step,” the exact E-step is this ED reference, and the contribution of the variational methods is to *generate* that same state variationally on a circuit rather than to load a classically-computed one.

#### C.2 Quantum technique common to both variational preparations

VQT and VarQITE share three ingredients, which we state once here so that the method-specific subsections can focus on what is genuinely different (how each reaches the thermal state and what it costs).

**Target and readout.** Both prepare, per sample $i$, the Gibbs state $\rho_i(s)$ on a $n_q = \lceil \log K \rceil$-qubit system register, and read out the responsibilities as the computational-basis populations $r_{ik} = \langle k|\, \rho_i(s) \,|k\rangle$. Both reduce to the exact classical softmax at the schedule endpoint $s{=}0$.

**Hardware-efficient brick-wall ansatz.** Both use the same family of parameterised circuit: a brick-wall of single-qubit $R_y, R_z$ rotations on every register qubit, followed per layer by a ladder/ring of CNOT entanglers. They differ only in what the ansatz acts on and how deep it is: VQT acts on the $n_q$-qubit system register at depth d, while VarQITE acts on a $2n_q$-qubit system–ancilla register at depth d (the extra register and depth are needed to entangle the purification across the system/ancilla boundary, Sec. C.4). At $\theta$=0 both ansätze reduce to the identity-block (rotations are the identity, CNOTs a basis permutation); a small Gaussian kick $\theta \sim \mathcal{N}(0, \varsigma^2)$, $\varsigma$=0.05, breaks the symmetry that would otherwise freeze the dynamics at the gradient-free maximally-mixed point, while staying within $\mathcal{O}(\varsigma^2)$ of it so no barren plateau is induced (the identity-block initialisation of Ref. [15]).

**Static Pauli-string Hamiltonian.** Both evaluate energies of $\hat{H}_i(s) = \sum_\mu h_\mu(s) P_\mu$ as Pauli-string expectations, $P_\mu \in \{\mathbb{I}, X, Y, Z\}^{\otimes n_q}$. The diagonal energy operator is expanded in $\{\mathbb{I}, Z\}$ strings by the discrete Walsh–Hadamard transform,

$$\mathrm{diag}(\mathbf{d}) = \sum_{\mathbf{z}\in\{0,1\}^{n_q}} c_\mathbf{z} \prod_{q:z_q=1} Z_q, \qquad c_\mathbf{z} = \frac{1}{K_\mathrm{pad}} \sum_k (-1)^{\mathbf{z}\cdot\mathbf{k}}\, d_k, \tag{S5}$$

which is $K_\mathrm{pad}$ strings whose operators are fixed across samples and schedule steps; only the coefficients $c_\mathbf{z}$ change. This static Pauli basis is what allows the entire $N$-sample E-step to be compiled once and vectorised. The transverse-field mixer contributes the $n_q$ strings $-\sum_a X_a$ (Appendix Appendix E treats the alternative cyclic-shift mixer).

#### C.3 Variational quantum thermalizer (VQT)

VQT [16] is the deployable preparation. It acts on the system register only, needs no ancilla, and turns the one genuinely hard part of thermal-state preparation—the entropy—into a closed-form classical quantity. In our setting it is applied per data point: at each variational iteration, every sample $i$ carries its own diagonal energy vector and VQT prepares a separate thermal state whose diagonal is read out as that sample's cluster responsibilities.

##### C.3.1 The free-energy variational principle

VQT represents the thermal state of $H_i(s)$ as a fixed brick-wall unitary $U(\theta)$ acting on a classical latent ensemble,

$$\rho_i(\theta, \phi) = U(\theta)\Big(\textstyle\sum_\mathbf{x} p_\phi(\mathbf{x})\ |\mathbf{x}\rangle\langle\mathbf{x}|\Big)U(\theta)^\dagger, \qquad p_\phi = \mathrm{softmax}(\phi), \tag{S6}$$

where $|\mathbf{x}\rangle$ ranges over the $K_\mathrm{pad}$ computational-basis states and $p_\phi$ is a classical distribution parameterised by $\phi \in \mathbb{R}^{K_\mathrm{pad}}$ through the softmax, which guarantees $p_\phi(\mathbf{x}) > 0$ and $\sum_\mathbf{x} p_\phi(\mathbf{x}) = 1$ without constraint. The variational free energy at temperature $T = 1/\beta$ is

$$\mathcal{F}_i(\theta, \phi) = \mathrm{Tr}\big[\rho_i(\theta, \phi)\, H_i(s)\big] - T\, S\big(\rho_i\big), \qquad S(\rho) = -\mathrm{Tr}(\rho \ln \rho). \tag{S7}$$

$\mathcal{F}_i$ is the Gibbs free energy and is stationary exactly when $\rho_i = e^{-\beta H_i(s)}/Z_i$, at which point $p_\phi$ equals the Gibbs eigenvalue spectrum and $U(\theta)$ rotates the latent into the Gibbs eigenbasis.

**The entropy is classical and exact.** Because $U(\theta)$ is unitary it leaves the spectrum of $\rho_i$ unchanged, so the eigenvalues of $\rho_i(\theta, \phi)$ are exactly the latent weights $p_\phi(\mathbf{x})$. The von Neumann entropy therefore collapses in closed form to the Shannon entropy of the latent,

$$S(\rho_i) = -\sum_{\mathbf{x}} p_\phi(\mathbf{x}) \ln p_\phi(\mathbf{x}), \tag{S8}$$

computed classically with no quantum estimation. This is the crux of why VQT is tractable: entropy is non-linear in $\rho$ and has no direct Pauli-expectation estimator—it is the obstruction that makes general thermal-state preparation hard—and Eq. (S8) removes it entirely. The energy term is a sum of Pauli expectations on a *static* basis (the Pauli decomposition of $H_i(s)$ does not depend on $\theta$), so the whole objective is measurable with a fixed set of observables.

##### C.3.2 The objective as implemented

We give the construction in full so that it can be reproduced. The input to the E-step for sample $i$ is the length-$K$ energy vector $D_i$; the output is the responsibility vector $r_i$. The objective is the free energy (S7) written on the latent basis,

$$\mathcal{L}(\theta, \phi) = \underbrace{\sum_{\mathbf{x}} p_\phi(\mathbf{x})\, E_{\mathbf{x}}(\theta; s)}_{\mathrm{Tr}[\rho_i H_i(s)]} + \underbrace{T \sum_{\mathbf{x}} p_\phi(\mathbf{x}) \ln p_\phi(\mathbf{x})}_{-T\, S(\rho_i)}, \qquad E_{\mathbf{x}}(\theta; s) = \langle \mathbf{x} |\, U(\theta)^\dagger H_i(s)\, U(\theta)\, | \mathbf{x} \rangle\,, \tag{S9}$$

and it is assembled and minimised in the following steps.

**Step 1: condition and pad the energies.** $D_i$ is shifted so its minimum is zero and, when non-degenerate, rescaled to a fixed range $[0, 4]$,

$$\tilde{d}_{ik} = 4\, \frac{D_{ik} - \min_{k'} D_{ik'}}{\max_{k'} D_{ik'} - \min_{k'} D_{ik'}}, \qquad k = 1, \ldots, K, \tag{S10}$$

(the shift is a global energy offset, invisible to the Gibbs state; the fixed range keeps $\beta\, \tilde{d}$ in a stable numerical window across cells). The vector is padded from $K$ to $K_{\mathrm{pad}} = 2^{n_q}$ by setting the $K_{\mathrm{pad}} - K$ padding entries to a large constant $\Lambda = 10^3$, so those levels receive weight $e^{-\beta\Lambda} \approx 0$ and do not participate; the padded vector is $\bar{d}_i \in \mathbb{R}^{K_{\mathrm{pad}}}$.

**Step 2: build the Hamiltonian in the Pauli basis.** The diagonal $\mathrm{diag}(\bar{d}_i)$ is expanded in Pauli-$Z$ strings by the Walsh–Hadamard transform: writing $\langle z, k \rangle$ for the bitwise dot product of the $n_q$-bit indices $z, k$ (MSB first), the coefficient of the $Z$-string labelled $z$ is

$$c_{iz}^{(Z)} = \frac{1}{K_{\mathrm{pad}}} \sum_{k=0}^{K_{\mathrm{pad}}-1} (-1)^{\langle z,k \rangle}\, \bar{d}_{ik}, \qquad \mathrm{diag}(\bar{d}_i) = \sum_z c_{iz}^{(Z)}\, Z_z, \tag{S11}$$

where $Z_z = \bigotimes_q Z_q^{z_q}$. Assembling the anneal path $H_i(s) = (1-s)\,\mathrm{diag}(\bar{d}_i) - s\, H_{\mathrm{mix}}$ then gives the full Pauli coefficient vector

$$c_i \;=\; \big[\, (1-s)\, c_{i,0}^{(Z)}, \ldots, (1-s)\, c_{i,K_{\mathrm{pad}}-1}^{(Z)}\,;\; +s, \ldots, +s \,\big], \tag{S12}$$

the first $K_{\mathrm{pad}}$ entries being the scaled diagonal coefficients and the last $n_q$ entries the transverse-field strings $\{X_q\}$ (for the cyclic-shift mixer these are instead the fixed coefficients of its dense Pauli decomposition, each multiplied by $-s$). Crucially this Pauli *basis* is static—it does not depend on $\theta$, $\phi$, or the sample—so it is built once and only the scalar coefficients $c_i$ change per sample and per anneal step.

**Step 3: evaluate all basis-state energies exactly.** For each computational basis state $|\mathbf{x}\rangle$, $\mathbf{x} = 0, \ldots, K_{\text{pad}} - 1$, the energy is the coefficient-weighted sum of Pauli expectations in the *rotated* state $U(\theta)\,|\mathbf{x}\rangle$,

$$E_{\mathbf{x}}(\theta; s) = \sum_p c_{i,p}\ \langle \mathbf{x}|\, U(\theta)^\dagger P_p\, U(\theta)\, |\mathbf{x}\rangle\,, \tag{S13}$$

and because $K_{\text{pad}} \leq 32$ in all our experiments we evaluate $E_{\mathbf{x}}$ for *all* $K_{\text{pad}}$ states by direct enumeration: one shallow-circuit preparation of $U(\theta)\,|\mathbf{x}\rangle$ per state, from which every $\langle P_p \rangle$ is read. No latent samples are drawn, so $\mathcal{L}$ in Eq. (S9) is computed with the exact $\{E_{\mathbf{x}}\}$ and $p_\phi = \text{softmax}(\phi)$.

**Step 4: gradients.** $\partial\mathcal{L}/\partial\theta$ uses the parameter-shift rule: each of the $|\theta|$ angles is evaluated at $\pm\pi/2$ shifts, so the quantum cost is $2|\theta|\,K_{\text{pad}}$ circuit evaluations per step. The $\phi$-gradient is closed form and adds *no* quantum work, since it reuses the already-measured $\{E_{\mathbf{x}}\}$: with $p_\phi = \text{softmax}(\phi)$ and $A_{\mathbf{x}} = E_{\mathbf{x}}(\theta; s) + T\big(\ln p_\phi(\mathbf{x}) + 1\big)$,

$$\frac{\partial \mathcal{L}}{\partial \phi_{\mathbf{x}}} = p_\phi(\mathbf{x})\Big(A_{\mathbf{x}} - \sum_{\mathbf{y}} p_\phi(\mathbf{y})\, A_{\mathbf{y}}\Big). \tag{S14}$$

(In our runs $\partial\mathcal{L}/\partial\theta$ is obtained by reverse-mode automatic differentiation through the same enumerated expectations, which is numerically equivalent to the parameter-shift estimator and cheaper in simulation; on hardware one substitutes the parameter-shift values.)

**Step 5: minimise.** $(\theta, \phi)$ are updated by Adam on the single objective (S9), with learning rates $(\eta_\theta, \eta_\phi) = (0.05, 0.1)$ and standard moments $(\beta_1, \beta_2, \varepsilon) = (0.9, 0.999, 10^{-8})$, for $n_{\text{steps}}$ iterations (updating both blocks jointly; an alternating schedule that updates $\theta$ and $\phi$ on odd/even steps is also supported). The loop is initialised warm from the previous outer iteration's $(\theta, \phi)$ when available, else at $\theta = 0$ (identity circuit) and $\phi = 0$ (uniform $p_\phi$); the $\theta = 0$, $s = 1$ start is the driver ground state and makes the E-step initialisation-independent. Every sample's inner loop is identical in structure, so the $N$ per-sample optimisations are vectorised into one batched, compiled kernel.

##### C.3.3 Readout

The prepared state's diagonal in the cluster basis gives the responsibilities directly. Since $\rho_i(\theta, \phi) = \sum_{\mathbf{x}} p_\phi(\mathbf{x}) U(\theta)\,|\mathbf{x}\rangle\langle\mathbf{x}|\, U(\theta)^\dagger$,

$$\big[\rho_i\big]_{kk} = \sum_{\mathbf{x}} p_\phi(\mathbf{x})\, \big|\langle k|U(\theta)|\mathbf{x}\rangle\big|^2, \tag{S15}$$

each term $|\langle k|U(\theta)|\mathbf{x}\rangle|^2$ being a single measured basis-state probability. The sample's responsibility vector is the $K$-block of this diagonal, clipped to be positive and renormalised, $r_{ik} = [\rho_i]_{kk} / \sum_{k'=1}^{K} [\rho_i]_{k'k'}$, so the padding levels are discarded at readout. These $r_{ik}$ are returned to the SVVS M-step, closing the variational loop.

##### C.3.4 Why it is hardware-ready

VQT is a single fixed circuit $U(\theta)$ on $n_q$ qubits: there is no ancilla register, no Hadamard-test auxiliary wire, and no imaginary-time trajectory to integrate. The latent $p_\phi$ and the gradient live classically; the only quantum operations are preparing $U(\theta)\,|x\rangle$ and measuring Pauli expectations. The same circuit can be run on a simulator and on hardware and the two outputs compared directly, which makes VQT the natural vehicle for simulator-versus-hardware diagnosis. Depth

$d$=3 suffices on the system-only register because $U(\theta)$ need only rotate the (already-classical) latent spectrum into the Gibbs eigenbasis, a shallower task than the cross-register entangling VarQITE must perform.

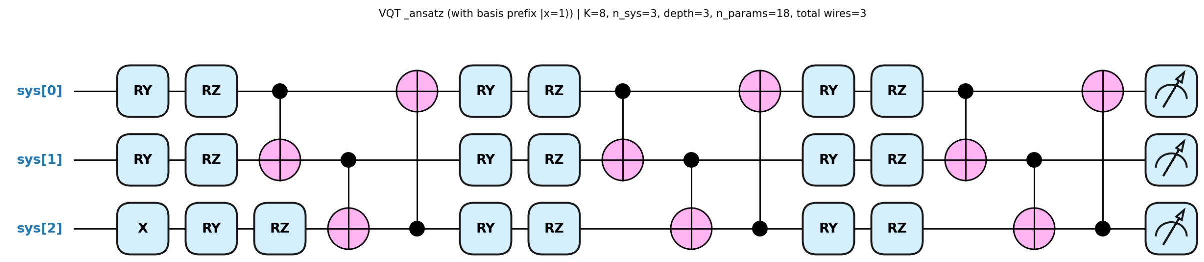

**Supplementary Figure S1**: Quantum circuit architecture of the VQT ansatz for a 8-cluster task. The schematic details the parameterization framework of the Variational Quantum Thermalization (VQT) ansatz tailored for a problem decomposed into (K = 8) clusters. The circuit utilizes ($n_{\text{sys}} =$ 3) system qubits to represent 8 distinct cluster states. An initial X gate is applied as a basis prefix to the third register to initialize the system into the specified reference state basis $|x = 1\rangle$. The ansatz is structured with a variational depth of d = 3. Each layer contains parameterized single-qubit rotations, RY and RZ, followed by a hardware-efficient network of controlled-NOT (CNOT) gates providing linear entanglement between adjacent registers. The entire architecture tracks a total of ($n_{\text{params}} = 18$) variational parameters prior to final readout measurements.

#### C.4 Variational imaginary-time evolution (VarQITE)

VarQITE [13, 14] is the more expensive anchor. It prepares the same Gibbs target by imaginary-time evolution of a purification, and its role here is to confirm that the variational preparations reproduce the exact thermal state (Sec. Appendix D); we therefore give its derivation concisely.

##### C.4.1 Imaginary-time evolution and the thermofield double

The Schrödinger equation in imaginary time $\tau$ reads $\partial_\tau |\psi(\tau)\rangle = -\hat{H}\,|\psi(\tau)\rangle$, with the non-unitary solution $|\psi(\tau)\rangle = e^{-\tau\hat{H}}\,|\psi(0)\rangle$. To realise a mixed Gibbs state on a device that produces pure states we use the thermofield-double (TFD) purification. Doubling the Hilbert space into system $S$ and ancilla $A$, each of dimension $K_{\text{pad}} = 2^{n_q}$,

$$|\text{TFD}(\beta)\rangle = \frac{1}{\sqrt{Z(\beta)}}\sum_k e^{-\beta E_k/2}\,|\phi_k\rangle_S \otimes |\phi_k\rangle_A\,, \qquad \hat{H}_S\,|\phi_k\rangle = E_k\,|\phi_k\rangle\,, \tag{S16}$$

has reduced state exactly the target Gibbs state,

$$\rho_S = \text{Tr}_A\big[\,|\text{TFD}(\beta)\rangle\langle\text{TFD}(\beta)|\,\big] = \frac{e^{-\beta\hat{H}_S}}{Z(\beta)}. \tag{S17}$$

At $\beta$=0 the TFD is the maximally entangled state $|\text{TFD}(0)\rangle = K_{\text{pad}}^{-1/2}\sum_k |k\rangle_S\,|k\rangle_A = \prod_q \text{CNOT}_{q\to n_q+q}\,\text{H}_q\,|0\rangle^{\otimes 2n_q}$, with reduced state $\mathbb{I}/K_{\text{pad}}$ (the $s_0$=1 uniform responsibilities). Reaching finite $\beta$ requires the non-unitary operator $e^{-\beta\hat{H}_S/2}$ on the system half, via $|\text{TFD}(\beta)\rangle \propto (e^{-\beta\hat{H}_S/2}\otimes\mathbb{I}_A)\,|\text{TFD}(0)\rangle$. A unitary circuit cannot apply this directly, so VarQITE approximates the trajectory variationally.

##### C.4.2 McLachlan's variational principle

Take a parameterised circuit $|\psi(\theta)\rangle = U(\theta)\,|0\rangle^{\otimes 2n_q}$ and differentiate along the manifold, $|\dot{\psi}\rangle = \sum_j \dot{\theta}_j\,|\partial_j\psi\rangle$. The exact imaginary-time flow satisfies $|\dot{\psi}\rangle = -(\hat{H} - \langle\hat{H}\rangle)\,|\psi\rangle$. The manifold cannot realise this exactly; McLachlan's principle [14, 13] chooses $\dot{\theta}$ to minimise the residual norm

$$\mathcal{L}(\dot{\theta}) = \Big\| \textstyle\sum_j \dot{\theta}_j\,|\partial_j\psi\rangle + (\hat{H} - \langle\hat{H}\rangle)\,|\psi\rangle \Big\|^2. \tag{S18}$$

The stationarity condition $\partial\mathcal{L}/\partial\dot{\theta}_i = 0$, using $\partial_j\langle\psi|\psi\rangle = 0$, gives the McLachlan equation

$$A(\theta)\,\dot{\theta} = -\,C(\theta), \qquad A_{ij} = \mathrm{Re}\big(\langle\partial_i\psi|\partial_j\psi\rangle - \langle\partial_i\psi|\psi\rangle\langle\psi|\partial_j\psi\rangle\big), \qquad C_i = \mathrm{Re}\,\langle\partial_i\psi|\hat{H}_i(s)|\psi\rangle. \tag{S19}$$

Here $A$ is the real part of the quantum Fisher information, i.e. the Fubini–Study metric on the variational manifold, and $C$ is the gradient of $\langle\hat{H}\rangle$; the update is natural-gradient descent on the energy. For a real-amplitude observable the identity $2\mathrm{Re}\langle\partial_i\psi|\hat{H}|\psi\rangle = \partial_i\langle\hat{H}\rangle$ reduces the McLachlan vector to half the energy gradient, $C_i = \frac{1}{2}\,\partial_i\langle\hat{H}\rangle_\theta$. When the ansatz is over-parameterised, $A$ is singular; we use Tikhonov regularisation $(A + \delta\mathbb{I})\dot{\theta} = -C$ with $\delta = 10^{-4}$, which keeps the solve stable and converges to the Moore–Penrose pseudoinverse as $\delta \to 0^+$.

##### C.4.3 Why the purification ansatz needs both registers

The one VarQITE-specific subtlety is why the ansatz must act on both halves of the purification rather than the system alone. A system-only ansatz $U(\theta) = U_S(\theta) \otimes \mathbb{I}_A$ fails: for the maximally entangled state,

$$\mathrm{Tr}_A\big[(U_S \otimes \mathbb{I}_A)\,|\mathrm{TFD}(0)\rangle\langle\mathrm{TFD}(0)|\,(U_S^\dagger \otimes \mathbb{I}_A)\big] = U_S\,\frac{\mathbb{I}}{K_{\mathrm{pad}}}\,U_S^\dagger = \frac{\mathbb{I}}{K_{\mathrm{pad}}}, \tag{S20}$$

so any unitary on one half leaves the reduced state of the other invariant and responsibilities stay pinned at uniform. The brick-wall ansatz of Sec. C.2 is therefore applied across the full $2n_q$-qubit register at depth $d$=3, with the per-layer entanglers including inter-half CNOTs that couple system qubit $q$ to ancilla qubit $q$; these inter-half entanglers are the load-bearing element that lets the reduced state leave the maximally mixed point, and they are why VarQITE needs the larger register and greater depth than VQT (Supplementary Figure S2). At $\theta$=0 the ansatz reduces to $|\mathrm{TFD}(0)\rangle$ and the McLachlan flow is started with the Gaussian kick of Sec. C.2. The McLachlan vector $C$ requires $\langle\hat{H}_S\rangle$, evaluated on the static Pauli basis of Sec. C.2.

#### C.5 Quantum-circuit resource comparison

The two variational preparations differ markedly in their circuit resources, and the difference is what makes VQT the more economical and the more hardware-ready of the two. We give the resource scalings in closed form and then tabulate the concrete counts at the cluster numbers used in this work.

The cost is driven by the register width and the variational circuit, and on both axes VQT is the leaner method. VQT acts on the system register only ($n_q$ qubits, no ancilla, no auxiliary wire), with $|\theta| = 2n_q d$ circuit angles at depth $d$=2 plus a classical latent of $K_{\mathrm{pad}} = 2^{n_q}$ probabilities $p_\phi$ whose entropy is evaluated in closed form (Eq. (**??**)) rather than on the device; its free-energy gradient is a single $O(|\theta|)$ pass because the entropy term is classical. VarQITE, by contrast, purifies the Gibbs state on a thermofield double, so it requires a system register of $n_q$ qubits, an equal-size ancilla register, and one auxiliary qubit for the Hadamard-test metric, for a total of $2n_q + 1$ qubits;

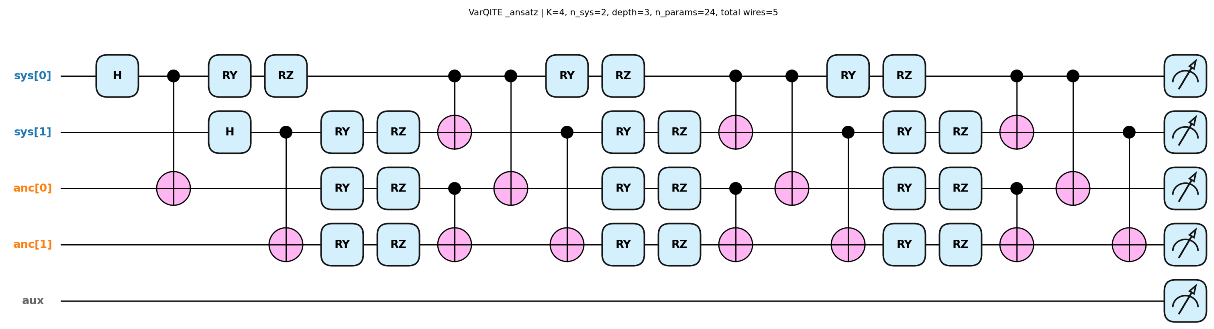

**Supplementary Figure S2**: Quantum circuit architecture of the VarQITE ansatz for a 4-cluster task. The diagram illustrates the parameterization structure for a variational quantum imaginary time evolution (VarQITE) simulation spanning (K = 4) target clusters. The architecture utilizes ($n_{\text{sys}} = 2$) system qubits, two ancilla qubits (anc[0], anc[1]), and one uncoupled auxiliary wire (aux), totaling 5 wires. Following initial entangling blocks, the ansatz scales via a hardware-efficient repeating structure of depth (d = 3). Each layer applies parameterized single-qubit rotations, RY and RZ, across all active system and ancilla qubits, followed by a hardware-efficient entangling network of controlled-NOT (CNOT) gates. The complete network maps a total of ($n_{\text{params}} = 24$) variational parameters prior to final projective measurements.

its brick-wall body carries two rotations $(R_y, R_z)$ on each of the $2n_q$ register qubits per layer at depth $d$=3, giving $|\theta| = 4n_q d$ angles, and each step must build the quantum Fisher-information metric $A$—an $O(|\theta|^2)$ object plus a linear solve. Table S8 gives the general formulas and Table S9 the concrete counts.

**Table S8:** Resource scaling of the three preparations of the quantum-coupled E-step, as a function of the cluster number through $n_q = \lceil \log K \rceil$. Circuit angles are the trainable rotation parameters $|\theta|$; VQT additionally optimises a classical latent of $K_{\text{pad}} = 2^{n_q}$ probabilities whose entropy is closed-form. The entangler-per-layer counts are read directly from the implemented brick-wall ansatz (VarQITE: system ring + ancilla ring + inter-half CNOTs, plus a fixed $n_q$-CNOT thermofield-double prefix; VQT: a single even/odd CNOT ladder with ring closure for $n_q \geq 3$).

| | qubits | circuit angles $\lvert\theta\rvert$ | entropy | per-step gradient |
|---|---|---|---|---|
| ED | $n_q$ | — | exact (ground truth) | — |
| VQT | $n_q$ | $2n_q d$ | exact, classical (Shannon) | $O(\lvert\theta\rvert)$ |
| VarQITE | $2n_q + 1$ | $4n_q d$ | not required (imag. time) | $O(\lvert\theta\rvert^2)$ (QFI metric + solve) |

### Appendix D Faithfulness of the simulation

All preparations are simulated as exact, noiseless statevectors, so any reported advantage is a property of the inference algorithm and not of a hardware or noise model. We verify faithfulness three independent ways.

**(1) Hamiltonian exactness.** The static Walsh–Hadamard $Z$-string expansion of Eq. (S11) reproduces the diagonal energy operator $\hat{H}_{E,i}$ to machine precision (maximum absolute error

**Table S9:** Concrete quantum-circuit resources at the cluster numbers used in this work, for the depths reported in the main text ($d$=3 for VarQITE on the system–ancilla register, $d$=2 for VQT on the system register). Total CNOTs counts the full depth-$d$ entangling circuit (VarQITE includes its fixed $n_q$-CNOT thermofield-double prefix).

| | | VQT ($d$=2) | | | | VarQITE ($d$=3) | | |
|---|---|---|---|---|---|---|---|---|
| $K$ | $n_q$ | qubits | angles | latent $p_\phi$ | total CNOTs | qubits | angles | total CNOTs |
| 2 | 1 | 1 | 4 | 2 | 0 | 3 | 12 | 4 |
| 3 | 2 | 2 | 8 | 4 | 2 | 5 | 24 | 14 |
| 4 | 2 | 2 | 8 | 4 | 2 | 5 | 24 | 14 |
| 8 | 3 | 3 | 12 | 8 | 6 | 7 | 36 | 30 |
| 16 | 4 | 4 | 16 | 16 | 8 | 9 | 48 | 40 |

Counts are exact for the implemented ansätze. The per-step gradient cost is $O(|\theta|)$ for VQT (free-energy pass, classical entropy) and $O(|\theta|^2)$ for VarQITE (quantum Fisher-information metric plus linear solve). VQT angles $= 2n_q d$; total CNOTs $= d\,[\,\lfloor n_q/2\rfloor + \lfloor (n_q-1)/2\rfloor + \mathbf{1}\{n_q \geq 3\}\,]$; classical latent $= 2^{n_q}$ probabilities. VarQITE angles $= 4n_q d$; total CNOTs $= n_q$ (TFD prefix) $+\,d\,[\,2(n_q-1) + 2\cdot\mathbf{1}\{n_q \geq 3\} + n_q\,]$. The clustering register used in the main experiments is $K$=3 ($n_q$=2) and the over-specified case $K$=4 ($n_q$=2); the $K \in \{8, 16\}$ rows indicate how the resources scale.

$< 10^{-12}$), so the simulated Hamiltonian is exactly.

**(2) Classical-limit recovery.** The automatic-differentiation quantum Fisher information used in VarQITE agrees with the analytically constructed metric, and at the schedule endpoint $s$=0 every preparation returns exactly the classical Boltzmann responsibilities; at the maximally mixed initialisation the prepared reduced state is the uniform distribution to $< 10^{-12}$.

**(3) Trace-distance agreement with the exact Gibbs state.** On the clustering register the VarQITE and VQT responsibilities reproduce the exact ED populations to within the integration tolerance. The informative check is the trace distance $T(\rho_{\text{var}}, \rho_{\text{exact}}) = \frac{1}{2}\|\rho_{\text{var}} - \rho_{\text{exact}}\|_1$, expected to be $\sim 10^{-2}$–$10^{-3}$ at moderate ansatz depth (*not* $10^{-7}$, since the finite manifold cannot contain the exact Gibbs state). Table S10 reports the depth dependence at a representative cell.

**Table S10:** VarQITE faithfulness at $K$=4, $\beta$=2, $s$=0.5, Gaussian-kick $\varsigma$=0.05, 40 McLachlan steps. Trace distance to the exact Gibbs state and the responsibility overlap $\sum_k \min(r_k^{\text{var}}, r_k^{\text{exact}})$.

| ansatz depth | steps | trace distance | overlap |
|---|---|---|---|
| 1 | 40 | 0.93 | 0.47 |
| 2 | 40 | 0.19 | 0.97 |
| 3 | 40 | 0.34 | 0.81 |
| 4 | 40 | 0.03 | 0.98 |

This single-cell sweep is the depth-selection diagnostic for VarQITE: every depth $\geq 2$ reaches high responsibility overlap ($\geq 0.81$), confirming the variational manifold contains a faithful approximation to the exact Gibbs state. The non-monotone trace distance at depth 3 reflects a known McLachlan optimisation pathology—the imaginary-time flow visits a poor saddle at this particular $(\beta, s)$ cell—not a failure of the depth; across the full annealing schedule depth $d$=3 is the robust choice and is the deployed VarQITE setting. VQT uses $d$=2 on its smaller system-only register (Sec. C.5), where the shallower circuit already converges. The responsibilities used downstream are read from the converged trajectory, and on the clustering register VarQITE and VQT agree with the exact ED populations to within the integration tolerance.

### Appendix E Robustness to the choice of mixing Hamiltonian

The original quantum-annealing variational Bayes (QAVB) formulation [10, 11] drives the E-step with a single-qudit cyclic-shift mixer $\hat{H}^{\mathrm{cyc}}_{\mathrm{mix}}$, the $K \times K$ matrix with $\langle k| \hat{H}^{\mathrm{cyc}}_{\mathrm{mix}} |k{\pm}1 \bmod K\rangle = 1$, whose ground state is the uniform superposition over assignments. In the main text we instead use the canonical multi-qubit transverse field $\hat{H}_{\mathrm{mix}} = -\sum_a X_a$, which is the standard quantum-annealing driver [12] and, being a sum of single-qubit operators, is the natural quantum object on the $\lceil \log K \rceil$-qubit assignment register. The transverse field also makes the falsifiable control of Appendix Appendix H exact: setting $\hat{H}_{\mathrm{mix}}{=}0$ recovers the strictly diagonal classical anneal.

To confirm that the reported clustering advantage is a property of the quantum-coupled E-step and not an artefact of the driver choice, we repeated the experiment at the reference cell ($N{=}400$, $S{=}5000$, $K{=}3$, $\sigma \approx 0.29$; over-specified truncation $K_{\mathrm{max}}{=}4$) with the cyclic-shift driver substituted for the transverse field and all other hyperparameters held fixed: identical annealing schedule, ansatz depth, optimisation budget, dataset, and seed sequence. Only the mixer changes. Table S11 reports the per-seed reliability of the VQT preparation under the two drivers.

**Table S11:** Per-seed reliability of the VQT quantum E-step at the reference cell ($\sigma \approx 0.29$, $K_{\mathrm{max}}{=}4$) under the two mixing Hamiltonians, with all other hyperparameters fixed. $P(\mathrm{ARI} \geq 0.4)$ is the success rate; the parallel-tempering baseline scores 0.00 on this metric at this cell (Appendix Appendix H). Both rows are full $R{=}100$ paired-seed runs on the identical dataset, differing only in the mixing Hamiltonian. The two drivers occupy the same regime—near-identical mean ARI (0.363 vs 0.364), both recover the true cluster count in $\approx 80\%$ of runs, and both clear parallel tempering—so the advantage is attributable to the off-diagonal coupling of the quantum E-step, not to the specific form of the mixer.

| mixing Hamiltonian | $R$ | mean ARI | median | best | $P(\mathrm{ARI}\geq 0.4)$ | $P(\hat{K}{=}3)$ |
|---|---|---|---|---|---|---|
| Transverse field $-\sum_a X_a$ (main text) | 100 | 0.363 | 0.384 | 0.676 | 0.47 | 0.89 |
| Cyclic shift (original QAVB) | 100 | 0.364 | 0.430 | 0.707 | 0.56 | 0.79 |

Both runs use the same $R{=}100$ paired seeds, so the comparison is seed-matched. The mean ARI is essentially identical across the two drivers (0.363 vs 0.364); the small differences in success rate (0.47 vs 0.56) and model-selection accuracy (0.89 vs 0.79) are within seed-to-seed variation, and we do not claim either driver is superior. The defensible conclusion is quantitative invariance: the advantage over parallel tempering and the model-selection power survive the change of mixer. The transverse field remains the reported driver for the physical and structural reasons above.

### Appendix F The free-energy barrier and the signal-to-noise regime

This appendix states the barrier identity used in the main text, gives its closed form for our generative design, defines the signal fraction $\sigma$, and reports the machine-checked verification. We separate what is known from what is new: the identity of Lemma 1 is established, and the decomposition of that identity into population signal and estimation noise, together with the diagnostic it yields, is the contribution.

**Generative design.** Cluster $k$ has Dirichlet concentration $\alpha_k \in \mathbb{R}^S_{>0}$ at three levels: a signal block with $\alpha_{\mathrm{sig}} = \alpha_{\mathrm{cross}} + (\alpha_{\mathrm{max}} - \alpha_{\mathrm{cross}}) \cdot \mathrm{sep}$ and $\alpha_{\mathrm{max}} = 1.8$, the other clusters' blocks at $\alpha_{\mathrm{cross}} = 0.20$, and background at $\alpha_{\mathrm{bg}} = 0.05$. Sample $i$ in cluster $k$ has $p_i \sim \mathrm{Dir}(\alpha_k)$ and $x_i \sim \mathrm{Multinomial}(n_i, p_i)$ with $n_i \sim$ NegBin of mean 8000; each entry is zeroed independently with probability $\zeta$. We write

$L = \lfloor Sf/K \rfloor$ for the number of informative taxa per cluster and $\rho := \alpha_{\text{sig}}/\alpha_{\text{cross}} \geq 1$ for the separation ratio. The reference configuration is $N = 400$, $S = 5000$, $K = 3$, separation 0.2, balanced classes, zero-inflation 0.80 and informative-taxon fraction 0.15, giving $\rho = 2.6$.

#### F.1 The barrier is a known information divergence

For a hard clustering $c$ the plug-in multinomial energy is

$$E(c) = -\sum_i \sum_s x_{is} \log \hat{p}_{c_i,s}, \qquad \hat{p}_{k,s} = \frac{\sum_{i:c_i=k} x_{is}}{\sum_{i:c_i=k} n_i}, \tag{S21}$$

and for clusters $a, b$ the merged-saddle barrier is the likelihood cost of forcing $\mathcal{A} \cup \mathcal{B}$ through one centre instead of two, $B = E_{\text{merged}} - E_{\text{split}}$.

**Lemma 1** (Barrier as a merge cost). *With $p_M = \lambda \hat{p}_a + (1-\lambda)\hat{p}_b$ and $\lambda = N_a^{\text{tot}}/N_{ab}^{\text{tot}}$,*

$$B = N_a^{\text{tot}}\, D_{\text{KL}}(\hat{p}_a \| p_M) + N_b^{\text{tot}}\, D_{\text{KL}}(\hat{p}_b \| p_M) = N_{ab}^{\text{tot}}\, \text{JS}_\lambda(\hat{p}_a, \hat{p}_b). \tag{S22}$$

This identity is established and we claim no novelty for it. It is the merge cost of the agglomerative Information Bottleneck, and equivalently the likelihood-ratio ($G$-test) statistic for homogeneity of two multinomial samples; the second equality is the definition of the $\lambda$-weighted Jensen–Shannon divergence [4]. We restate the derivation in our notation because the decomposition in Section F.3 depends on this form.

*Proof.* Third clusters contribute identically to $E_{\text{merged}}$ and $E_{\text{split}}$ and cancel, so only $a$ and $b$ remain. For cluster $a$, $E^{(a)}_{\text{merged}} - E^{(a)}_{\text{split}} = \sum_s \left(\sum_{i\in\mathcal{A}} x_{is}\right) \log(\hat{p}_{a,s}/p_{M,s})$, and the MLE identity $\sum_{i\in\mathcal{A}} x_{is} = N_a^{\text{tot}} \hat{p}_{a,s}$ turns this into $N_a^{\text{tot}} D_{\text{KL}}(\hat{p}_a \| p_M)$. The same holds for $b$; substituting $p_M$ and expanding $H(p_M)$ gives the entropy-gap form. □

**Remark 1** (What $B$ measures). *Reading $B$ as a merge cost determines its interpretation. A merge cost is large exactly when the two clusters are well separated, so a tall barrier marks an* easy *instance. Barrier height therefore cannot order inference difficulty, which is what motivates the decomposition below.*

#### F.2 Closed form for the population signal

**Lemma 2** (Closed-form population signal). *For the symmetric block design,*

$$\text{JS}_{1/2}(\bar{p}_a, \bar{p}_b) = \frac{L\,\alpha_{\text{cross}}}{\alpha_0}\,\phi(\rho), \qquad \phi(\rho) = \rho \log \frac{2\rho}{\rho+1} + \log \frac{2}{\rho+1}, \tag{S23}$$

*with $\alpha_0 = L\alpha_{\text{sig}} + (K-1)L\alpha_{\text{cross}} + (S - Sf)\alpha_{\text{bg}}$. The function $\phi$ satisfies $\phi(1) = 0$ and is strictly increasing and convex on $[1, \infty)$. The signal barrier is $B_{\text{signal}} = N_{ab}^{\text{tot}}\, \text{JS}_{1/2}(\bar{p}_a, \bar{p}_b)$.*

*Proof.* Two clusters differ only on their $2L$ informative taxa, where the population profiles take values in ratio $\rho$; every other taxon contributes zero. The per-feature symmetric divergence is $\frac{m}{2}\phi(\rho)$ with $m = \alpha_{\text{cross}}/\alpha_0$, and summing over $2L$ taxa gives Eq. (S23). Monotonicity follows from $\phi'(\rho) = \log \frac{2\rho}{\rho+1} > 0$ for $\rho > 1$. □

**Remark 2** (Verification). *At $\rho = 2.6$ the closed form gives $\phi = 0.3683$ and $\text{JS}_{1/2} = 4.162 \times 10^{-2}$, matching direct numerical evaluation of the population divergence to four significant figures.*

##### F.3 Signal fraction

Splitting Eq. (S22) at its population value gives

$$B = B_{\text{signal}} + B_{\text{noise}}, \qquad B_{\text{signal}} = N_{ab}^{\text{tot}}\,\mathrm{JS}_\lambda(\bar{p}_a, \bar{p}_b), \qquad B_{\text{noise}} := B - B_{\text{signal}}, \tag{S24}$$

which is an identity by construction. Its content lies in the interpretation of the two terms: $B_{\text{signal}}$ is the separation between the populations, and $B_{\text{noise}}$ is the excess divergence contributed by estimating high-dimensional sparse profiles from finitely many counts. Plug-in divergence estimators are biased upward in this regime, so $\mathbb{E}[B_{\text{noise}}] \geq 0$; we state this in expectation because that is what the argument establishes, and we verified that $B_{\text{noise}} > 0$ in every cell of the regime sweep.

**Definition S1** (Signal fraction). *$\sigma := B_{\text{signal}}/B$, with $\mathrm{SNR} = \sigma/(1-\sigma) = B_{\text{signal}}/B_{\text{noise}}$. For $K > 2$ the difficulty-ordering quantity is the worst-pair value $\sigma_{\min} = \min_{a,b}\sigma_{ab}$ of Section F.5; in the symmetric design all pairs are equivalent and $\sigma_{\min} = \sigma$. Whenever $B_{\text{noise}} > 0$, which holds throughout our sweep, $\sigma \in [0, 1)$.*

**Observation 1** ($\sigma$ and the spurious-optimum gap). *Writing $\delta$ for the energy gap between the correct clustering and the nearest spurious optimum, we observe $\delta$ close to $B_{\text{signal}}$, and hence $B/\delta$ close to $1/\sigma$.*

##### F.4 Why $\sigma$ rather than $B$ orders difficulty

Deterministic annealing crystallises a split at a temperature set by the per-count signal, not by the total divergence [6, 7], $T_{\text{split}} \propto B_{\text{signal}}/N_{ab}^{\text{tot}}$. Applying the same relation to the noise component gives $T_{\text{noise}}/T_{\text{signal}} = B_{\text{noise}}/B_{\text{signal}} = (1-\sigma)/\sigma$, so whenever $\sigma < 0.5$ the noise-induced split crystallises at the higher temperature and a cooling schedule reaches it first. This accounts for the observation in the main text that deterministic annealing reaches the same partition as greedy ascent in every cell tested, and it predicts that widening the temperature range cannot repair this. We borrow the relation instead of deriving it for this model, and treat the consequence as a prediction that the sweep confirms.

##### F.5 Generalisation to $K$ clusters

A $K$-cluster solution has $\binom{K}{2}$ pairwise barriers, and difficulty is a property of that landscape. Lemmas 1 and 2 are the two-cluster case of results that hold for every pair, with $N_{ab}^{\text{tot}}$ in place of the global count and $\alpha_0$ and $L$ now functions of $K$.

**Lemma 3** (Signal dilution). *Writing $L \approx Sf/K$, the normaliser $\alpha_0(K) = \frac{Sf}{K}(\alpha_{\text{sig}} - \alpha_{\text{cross}}) + Sf\,\alpha_{\text{cross}} + (S - Sf)\alpha_{\text{bg}}$ approaches a constant as K grows, so $L/\alpha_0(K) \sim 1/K$ and each additional cluster shrinks every pair's distinguishing signal.*

**Remark 3** (Verification). *Machine-checked on the symmetric design: $K = 3$ gives $L = 250$, $\alpha_0 = 442.5$, $\mathrm{JS}_{1/2} = 0.04162$; $K = 6$ gives $L = 125$, $\alpha_0 = 402.5$, $\mathrm{JS}_{1/2} = 0.02288$. The ratio 0.550 is the $1/K$ law with the mild $\alpha_0$-saturation correction.*

Two channels lower $\sigma_{\min}$ as $K$ grows: the per-pair signal falls as $1/K$, and with roughly $N/K$ samples per cluster the finite-sample divergence bias rises. Recovery additionally faces a multiplicity of harmful spurious optima, from pairwise and three-way merges and label-swap confusions, that grows at least quadratically in $K$.

**Remark 4** (Scope). *Lemma 1 is established in the literature and restated here; Lemma 2 and Lemma 3 are exact for our design and machine-verified. Eq.* (S24) *is an identity by construction and the non-negativity of $B_{\text{noise}}$ holds in expectation. We make no closed-form claim for the single-run success probability of the quantum E-step at low $\sigma$: recovery there is a multi-well competition governed by the signal gap, the noise ruggedness and the multiplicity of spurious optima, and we characterise it by the empirical reliability curves rather than by a formula.*

### Appendix G Annealing schedule and stopping rule

A quantum-coupled E-step raises a legitimate question about what the algorithm returns. If a fit stopped while the mixer was still active, its output would not be a fixed point of the variational objective. Two properties of the procedure exclude this by construction, and we verify both empirically.

**Notation.** Let $t = 1, \ldots, T_{\max}$ index the outer variational iterations ($T_{\max} = 400$). At iteration $t$ the mixer strength and inverse temperature are set by the two-phase schedule

$$s_t = s_0 \max\Big(1 - \tfrac{t}{\tau_1},\, 0\Big), \qquad \beta_t = \begin{cases} \beta_0, & t \le \tau_1, \\ 1 + (\beta_0 - 1)\frac{\tau_2 - t}{\tau_2 - \tau_1}, & \tau_1 < t \le \tau_2, \\ 1, & t > \tau_2, \end{cases}$$

with $s_0 = 1$, $\beta_0 = 30$, $\tau_1 = 100$, $\tau_2 = 230$. The E-step branches on $s_t$: for $s_t > 0$ it prepares the annealed state $\rho \propto e^{-\beta_t H_S(s_t)}$ and reads the responsibilities off its diagonal; for $s_t = 0$ it is the ordinary $\beta_t$-scaled softmax of the classical variational update. Because $s_t = 0$ for every $t \ge \tau_1$, the mixer is off after iteration $\tau_1$ under every schedule; there is no setting in which it remains active.

**The stopping test cannot fire during the quantum phase.** The fit stops when the free-energy test declares convergence or when the iteration budget $T_{\max}$ is reached. The convergence test is evaluated only when three conditions hold simultaneously: (i) $t > \tau_2 = 230$; (ii) at least one pruning step has occurred; and (iii) at least three free-energy values are available (the free energy is recorded every ten iterations). Pruning itself is forced to begin no earlier than $\tau_1 + \Delta_{\text{prune}}$ (here $\Delta_{\text{prune}} = 5$), so the first prune, and hence the earliest admissible convergence, occurs strictly after the mixer has switched off. Condition (i) is the binding one: the earliest possible stop is iteration 231, which is 131 iterations after $s_t$ has reached zero. Every fit therefore ends with a long run of the unmodified classical update.

**Verification procedure.** Although the arguments above are structural, we verify them empirically (not by schedule inspection) for all three preparations—ED, VarQITE, and VQT—since any annealed E-step could in principle stop before reaching the classical phase. For each fit we record: the terminal outer-iteration index $t^\star$; the mixer strength $s_{t^\star}$; the indicator $\text{reached}_{s=0} = [t^\star \ge \tau_1 \wedge s_{t^\star} = 0]$; whether the free-energy convergence test fired (vs. hitting the budget); and the number of terminal classical iterations $t^\star - \tau_1$ after $s_t$ reached zero. The failure mode of concern is $t^\star < \tau_1$ (equivalently $\text{reached}_{s=0} = \mathsf{false}$). We run each preparation at three difficulty cells ($\sigma \in \{0.29, 0.43, 0.53\}$) with $R = 30$ random initialisations per cell (90 fits per method).

For all preparations and all 90 fits, runs entered the classical phase before stopping: $t^\star = 240$ in every case, at least 140 iterations beyond $\tau_1 = 100$. Thus $\text{reached}_{s=0} = \mathsf{true}$ for all 270 fits and no run stopped with an active mixer. VarQITE and VQT also terminate on the free-energy test more

| preparation | reached $s = 0$ | earliest stop $t^\star$ | terminal classical iters | free-energy test / budget |
|---|---|---|---|---|
| ED (exact) | 90/90 | 240 | $\geq 140$ | 39/51 |
| VarQITE | 90/90 | 240 | $\geq 140$ | 87/3 |
| VQT | 90/90 | 240 | $\geq 140$ | 89/1 |

often (87/90 and 89/90) than ED (39/90), consistent with slightly softer responsibilities yielding smoother free-energy plateaus and more frequent three-check certification within $T_{\max}$.

Budget-terminated runs (mainly ED at harder cells, where 22–25/30 hit the budget) are nevertheless at fixed points: in every inspected case, the relative free-energy change over the final interval—and over each of the last three intervals used by the test—was zero to numerical precision, well below $10^{-4}$. Non-firing is therefore a timing artifact: free energy is logged every ten iterations and certification requires three consecutive sub-tolerance intervals after both $t > \tau_2$ and the first prune, which some slowly settling runs may not satisfy before $T_{\max}$ despite having already plateaued. The budget thus stops runs that have ceased descending, not runs still evolving.

**Interpretation.** The quantum-coupled phase occupies the first $\tau_1$ iterations and selects which basin the fit enters. The final 140 or more iterations, which fix the reported responsibilities, are ordinary variational Bayes at $s = 0$, and this holds identically for the exact, VarQITE, and VQT preparations. QBayMic is therefore variational inference whose basin is chosen by an annealed quantum-coupled phase, and whose output is a fixed point of the same objective the classical arms optimise; the quantum coupling changes which optimum is reached, not what counts as a solution.

### Appendix H Controls and baselines

To attribute any advantage correctly we compare against three classical methods sharing the DMM–SVVS M-step and the evaluation protocol identically.

**Greedy variational inference (VB).** At $s$=0; the standard mean-field baseline.

**Deterministic-annealing VB (DAVB).** A classical control sharing the identical temperature schedule but with the mixing Hamiltonian removed ($\hat{H}_{\text{mix}}$=0). This isolates whether any gain is due to annealing alone or to the quantum mixer: DAVB anneals exactly as the quantum methods do, but its E-step remains strictly diagonal throughout, so any quantum advantage over DAVB cannot be a temperature-schedule effect.

**Parallel tempering (PT).** A replica-exchange scheme over an $M$-rung geometric inverse-temperature ladder whose cold replica ($\beta$=1) is exactly greedy VB. Each replica anneals through the same temperature-scaled E-step the quantum methods use, and adjacent rungs exchange configurations by a Metropolis swap on the common $\beta$=1 energy [17]. To make PT a strong rather than a straw-man baseline we add hot-replica reseeding from fresh basins (the mean-field analogue of a Monte-Carlo long jump) and gate swaps to equal cluster count. The ladder is tuned to a 20–40% swap-acceptance window. It tests whether the advantage is genuinely quantum (off-diagonal coupling) rather than merely better classical annealing. At matched compute (400 energy evaluations per method) and the reference cell with $K_{\max}$=4, PT reaches the success threshold in 0/100 seeds, while the quantum E-steps clear it in 47% (VarQITE/VQT) and 64% (ED); the paired effect sizes are $\Delta$ARI = +0.355 (VarQITE/VQT, 95% CI $[+0.317, +0.390]$, Wilcoxon $p = 5.6 \times 10^{-18}$)

and $\Delta$ARI $= +0.471$ (ED, 95% CI $[+0.425, +0.514]$, $p = 3.9 \times 10^{-18}$). The reliability advantage therefore survives parallel tempering.

#### Data and code availability

All quantum E-steps are simulated as exact, noiseless statevectors using PennyLane [18]. Code implementing the three state preparations (exact diagonalization, VarQITE, VQT), the Dirichlet–multinomial synthetic data generator, the parallel-tempering baseline, the barrier/$\sigma$ calculator, and the paired evaluation harness, together with the random seeds, the shared hyperparameter grid, and the pre-registered evaluation protocol, are available at `https://github.com/tungtokyo1108/Quantum_Bayes`.

#### References

[1] T. Dang, K. Kumaishi, E. Usui, S. Kobori, T. Sato, Y. Toda, Y. Yamasaki, H. Tsujimoto, Y. Ichihashi, and H. Iwata, *Stochastic variational variable selection for high-dimensional microbiome data*, Microbiome **10**, 236 (2022).

[2] I. Holmes, K. Harris, and C. Quince, *Dirichlet multinomial mixtures: generative models for microbial metagenomics*, PLoS ONE **7**(2), e30126 (2012).

[3] D. M. Blei, A. Kucukelbir, and J. D. McAuliffe, *Variational inference: a review for statisticians*, J. Am. Stat. Assoc. **112**(518), 859–877 (2017).

[4] J. Lin, *Divergence measures based on the Shannon entropy*, IEEE Trans. Inf. Theory **37**(1), 145–151 (1991).

[5] T. M. Cover and J. A. Thomas, *Elements of Information Theory*, 2nd ed., Wiley, Hoboken (2006).

[6] K. Rose, E. Gurewitz, and G. C. Fox, *Statistical mechanics and phase transitions in clustering*, Phys. Rev. Lett. **65**(8), 945–948 (1990).

[7] K. Rose, *Deterministic annealing for clustering, compression, classification, regression, and related optimization problems*, Proc. IEEE **86**(11), 2210–2239 (1998).

[8] A. Decelle, F. Krzakala, C. Moore, and L. Zdeborová, *Asymptotic analysis of the stochastic block model for modular networks and its algorithmic applications*, Phys. Rev. E **84**, 066106 (2011).

[9] L. Zdeborová and F. Krzakala, *Statistical physics of inference: thresholds and algorithms*, Adv. Phys. **65**(5), 453–552 (2016).

[10] H. Miyahara and V. Roychowdhury, *Quantum advantage in variational Bayes inference*, Proc. Natl. Acad. Sci. USA **120**(31), e2212660120 (2023).

[11] I. Sato, K. Kurihara, S. Tanaka, H. Nakagawa, and S. Miyashita, *Quantum annealing for variational Bayes inference*, in *Proc. 25th Conf. on Uncertainty in Artificial Intelligence (UAI)*, pp. 479–486 (2009); arXiv:0905.3528.

[12] T. Kadowaki and H. Nishimori, *Quantum annealing in the transverse Ising model*, Phys. Rev. E **58**, 5355–5363 (1998).

[13] S. McArdle, T. Jones, S. Endo, Y. Li, S. C. Benjamin, and X. Yuan, *Variational ansatz-based quantum simulation of imaginary time evolution*, npj Quantum Inf. **5**, 75 (2019).

[14] X. Yuan, S. Endo, Q. Zhao, Y. Li, and S. C. Benjamin, *Theory of variational quantum simulation*, Quantum **3**, 191 (2019).

[15] E. Grant, L. Wossnig, M. Ostaszewski, and M. Benedetti, *An initialization strategy for addressing barren plateaus in parametrized quantum circuits*, Quantum **3**, 214 (2019).

[16] G. Verdon, J. Marks, S. Nanda, S. Leichenauer, and J. Hidary, *Quantum Hamiltonian-based models and the variational quantum thermalizer algorithm*, arXiv:1910.02071 (2019).

[17] D. J. Earl and M. W. Deem, *Parallel tempering: theory, applications, and new perspectives*, Phys. Chem. Chem. Phys. **7**, 3910–3916 (2005).

[18] V. Bergholm et al., *PennyLane: automatic differentiation of hybrid quantum-classical computations*, arXiv:1811.04968 (2018).
